# Computational design of serine hydrolases

**DOI:** 10.1101/2024.08.29.610411

**Authors:** Anna Lauko, Samuel J. Pellock, Ivan Anischanka, Kiera H. Sumida, David Juergens, Woody Ahern, Alex Shida, Andrew Hunt, Indrek Kalvet, Christoffer Norn, Ian R. Humphreys, Cooper Jamieson, Alex Kang, Evans Brackenbrough, Asim K. Bera, Banumathi Sankaran, K. N. Houk, David Baker

## Abstract

Enzymes that proceed through multistep reaction mechanisms often utilize complex, polar active sites positioned with sub-angstrom precision to mediate distinct chemical steps, which makes their de novo construction extremely challenging. We sought to overcome this challenge using the classic catalytic triad and oxyanion hole of serine hydrolases as a model system. We used RFdiffusion^1^ to generate proteins housing catalytic sites of increasing complexity and varying geometry, and a newly developed ensemble generation method called ChemNet to assess active site geometry and preorganization at each step of the reaction. Experimental characterization revealed novel serine hydrolases that catalyze ester hydrolysis with catalytic efficiencies (*k*_*cat*_/*K*_*m*_) up to 3.8 × 10^3^ M^-1^ s^-1^, closely match the design models (Cα RMSDs < 1 Å), and have folds distinct from natural serine hydrolases. In silico selection of designs based on active site preorganization across the reaction coordinate considerably increased success rates, enabling identification of new catalysts in screens of as few as 20 designs. Our de novo buildup approach provides insight into the geometric determinants of catalysis that complements what can be obtained from structural and mutational studies of native enzymes (in which catalytic group geometry and active site makeup cannot be so systematically varied), and provides a roadmap for the design of industrially relevant serine hydrolases and, more generally, for designing complex enzymes that catalyze multi-step transformations.

## Main Text

Enzymes are exquisite catalysts that dramatically accelerate reaction rates in mild aqueous conditions. The ability to construct enzymes catalyzing arbitrary chemical reactions would have enormous utility across a wide range of applications, and hence, enzyme design has been a long-standing goal of computational protein design^2^. De novo enzyme design has generally started from a specification of arrangements of catalytic residues around the reaction transition state (a theozyme), and sought to identify placements of this active site in pre-existing scaffolds^3–8^. Fixed backbone scaffolds restrict how accurately the catalytic geometry can be realized, and this has likely limited the activities of many designed enzymes to date prior to optimization by laboratory evolution. A further challenge of enzyme design is the preorganization of the active site with atomic accuracy. Achieving preorganization is especially difficult for multistep reaction mechanisms, because the enzyme must preferentially stabilize multiple transition states and intermediates. Existing methods to evaluate design preorganization in silico^8–12^ are limited by low accuracy or computational cost and are typically only applied to one reaction state. To enable the accurate design of multistep enzymes, new methods are needed for both the generation of protein backbones tailored specifically to a given active site and assessment of their structural compatibility throughout the catalytic cycle.

We reasoned that advances in deep learning for protein design and structure prediction could be used to design proteins from scratch to scaffold a given active site and assess compatibility across a proposed reaction coordinate. Recent advances in scaffolding functional sites with RFdiffusion have yielded improved in silico and experimental success rates across a range of design tasks^1,13^; we aimed to use the same approach to generate enzymes starting from geometric descriptions of an active site (Fig. 1A). To assess preorganization and functional interactions in each step of the catalytic cycle, we sought to leverage advances in deep learning-based prediction of protein-small molecule complexes by modeling structural ensembles of catalytic intermediates (Fig. 1B).

**Figure 1.**
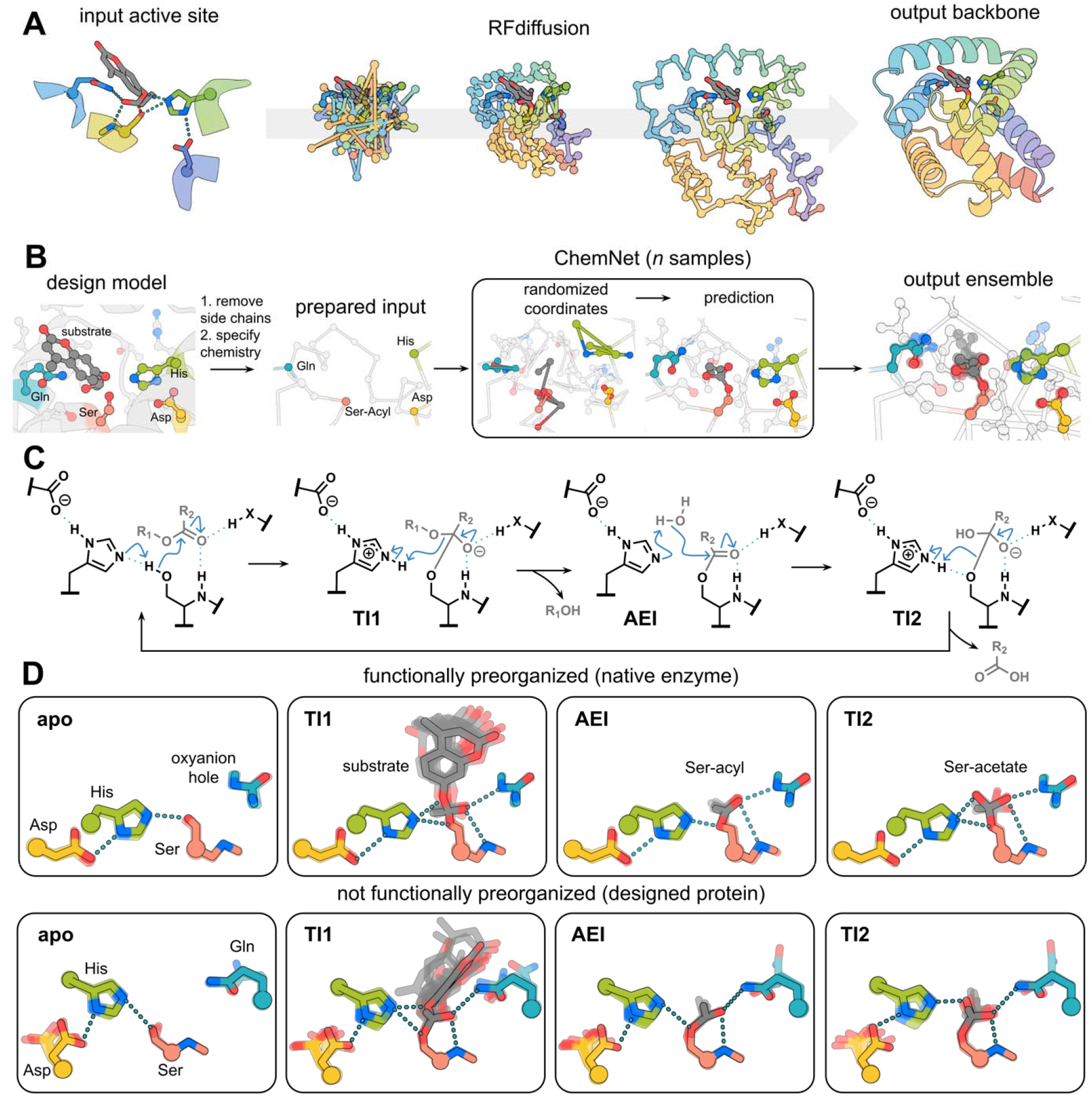
Design methods. **(A)** Active site specific backbone generation with RFdiffusion. Given the geometry of a possible active site configuration, RFdiffusion denoising trajectories generate backbone coordinates which scaffold the site. **(B)** Generation of active site ensembles with ChemNet. The coordinates of the sidechains around the active site and any bound small molecule for the step in the reaction being considered are randomized, and n samples are carried out to generate an ensemble of predictions. **(C)** Mechanism of ester hydrolysis by serine hydrolases. **(D)** Chemnet ensembles for distinct states along the reaction coordinate for hydrolysis of 4MU-Ac for a native serine hydrolase (top, PDB: 1IVY) and a designed serine hydrolase (bottom, josie).

Ester hydrolysis has served as a model reaction for computational enzyme design for decades^14–19^, and the catalytic triad and oxyanion hole of natural serine hydrolases utilize one of the most extensively studied enzymatic mechanisms to catalyze this reaction^20–27^. The catalytic cycle can be divided into four steps (Fig. 1C). First, the substrate binds to the apoenzyme (apo) and the catalytic serine, deprotonated by the catalytic histidine, attacks the carbonyl carbon of the ester to form the first tetrahedral intermediate (TI1). Second, the catalytic histidine protonates the leaving group oxygen promoting its departure, leaving the active site serine covalently linked to the acyl group of the substrate (acyl-enzyme intermediate, AEI). Third, the histidine deprotonates a water molecule, which attacks the AEI to generate a second tetrahedral intermediate (TI2). Finally, this intermediate is resolved by histidine-mediated protonation of serine and release of the acyl group, reconstituting the free enzyme and completing the catalytic cycle. Throughout, negatively charged transition states and intermediates are stabilized by a pair of hydrogen bond donors that constitute the oxyanion hole. Perturbation of the histidine pK_a_, which tunes its acid/base function, is mediated by interaction with aspartate or glutamate, the final residue in the triad^28–30^.

Despite extensive structural, mutational, and computational characterization of native serine hydrolases^31–34^, de novo design efforts that have attempted to employ this mechanism have been largely unsuccessful, yielding proteins that harbor activated serines and cysteines but fail to catalyze turnover^7,8^. We initially speculated that increasing scaffold diversity would help identify backbones that more accurately reconstruct the desired active site; and we carried out a preliminary design campaign searching for placements of a serine hydrolase active site in a library of deep-learning generated hallucinated NTF2 scaffolds that previously yielded catalysts for a luciferase reaction^35^. As in previous studies, experimental characterization of the resulting designs revealed activated serines but no catalytic turnover on activated ester substrates, despite a close match between the experimental and designed structures (Fig. S1), suggesting that key features important for catalysis were missing.

### Assessing reaction path compatibility with ChemNet

We set out to understand why these and earlier computational designs failed to catalyze ester hydrolysis and hypothesized that modeling states across the complete reaction coordinate could be critical for assessing the ability of a design to achieve catalytic turnover. To model the extent to which a designed enzyme can form each of the key states along the reaction cycle and to assess the preorganization of the active site residues in the desired catalytic geometries, we developed a deep neural network that, given (1) the backbone coordinates of a small molecule binding pocket or active site, (2) the identities of the amino acid residues at each position, and (3) the chemical structures of bound small molecules (but not their positions), generates the full atomic coordinates of the binding site, comprising both protein sidechains and small molecules. We trained this network, called ChemNet, on protein-small molecule complexes in the PDB by randomizing the atomic coordinates of sidechains and small molecules within spherical regions with up to 600 heavy atoms, and seeking to minimize a loss function assessing the recapitulation of the atomic coordinates within the region. ChemNet rebuilds regions within native structures with an average RMSD of 1.1 L. ChemNet is stochastic, and repeated runs from different random seeds yield an ensemble of models for the rebuilt region.

We used ChemNet to generate structural ensembles for each of the four reaction steps for a set of native and previously designed serine hydrolases. These calculations showed that native hydrolases are considerably more preorganized than previous designed systems (Fig. 1D, Fig. S2). In native systems, the catalytic residues at each step sample a very limited number of conformations in which all key hydrogen bonding interactions are maintained, but in designed systems there can often be wide variations in the ensembles at multiple steps. Since the reaction rate should be proportional to the fraction of the enzyme in the active state, the lack of preorganization of the designed active sites is expected to compromise catalysis. To quantify the extent of active site formation in the ChemNet ensembles, we compute the frequency of formation of key interactions between the catalytic functional groups and reaction intermediates over each step of the reaction (see Methods).

### Design and characterization of serine hydrolases

We next set out to design proteins with active sites of increasing complexity, using RFdiffusion to scaffold serine hydrolase active site motifs and ChemNet to assess their preorganization in each step of the reaction (Fig. 2A,B). We designed catalysts for the hydrolysis of 4-methylumbelliferone (4MU) esters (Fig. 2C) that fluoresce upon hydrolysis. To generate backbones to scaffold the catalytic machinery, we placed the catalytic sidechains around the substrate and starting from the backbone N, Cα, and C atoms of these key residues and their adjacent neighbors (i.e. a contiguous three-residue segment), used RFdiffusion to build up backbones, starting from random noise, which have coordinates that exactly match the input motif and also form a binding pocket for the substrate (see Methods). To drive folding to the designed state, and to make favorable interactions with the substrate and active site residues, LigandMPNN^36^ was used to design the sequence. Rosetta FastRelax^37^ was used to refine the protein backbone and ligand pose, and the sequence was again designed with LigandMPNN with the new backbone as input^38^. Following several iterations between LigandMPNN and FastRelax, the structures of the designs were predicted with AlphaFold2 (AF2)^39^, and designs for which all catalytic residue Cα’s were positioned within 1.0 Å of the design models were selected for experimental characterization^39^ (see Methods for additional details of computational design).

**Figure 2.**
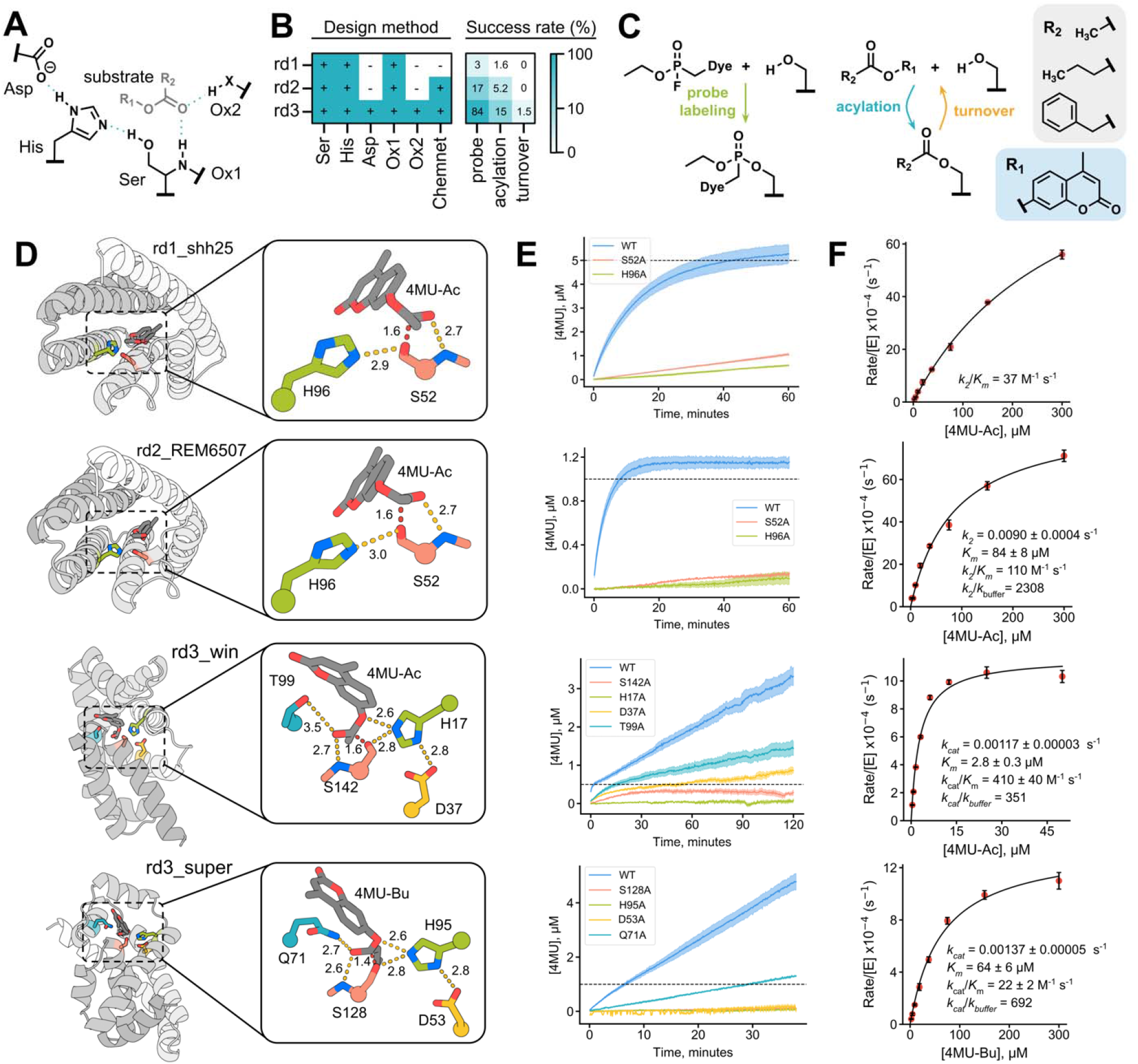
Functional characterization of designed serine hydrolases. **(A)** Chemical schematic of a serine hydrolase active site. **(B)** Summary of design method and experimental success rate for probe labeling, single turnover acylation, and catalytic turnover for each design round. **(C)** Chemical schematic depicting probe labeling, acylation, and catalytic turnover. **(D)** Fold (left) and active site (right) of serine hydrolase design models. **(E)** Reaction progress curves for the parent design and catalytic residue knockouts. Dashed line represents the enzyme concentration. **(F)** Michaelis-Menten plots derived from initial (rd1, rd2) or steady state velocities (rd3).

In the first two rounds of design, we built relatively simple active sites consisting of Ser-His dyads with a single oxyanion hole contact from the backbone amide of the serine (Fig. 2A,B), and explicitly evaluated the utility of ChemNet to select designs for experimental characterization; round 1 designs were filtered with AF2 alone, while round 2 designs that passed the AF2 filter were selected for experimental screening if ChemNet ensembles of the apo state indicated the key Ser-His hydrogen bond was formed (see Methods). Only 1.6% of round 2 designs passing AF2 filtering were predicted to be preorganized by ChemNet. For experimental testing, we obtained synthetic genes encoding 129 and 192 designs for rounds 1 and 2, respectively, for *E. coli* overexpression and screening.

We used a fluorophosphonate (FP) activity-based probe and fluorescent 4MU-acetate (4MU-Ac) and 4MU-butyrate (4MU-Bu) ester substrates to identify designs with activated serines and esterase activity, respectively (Fig. 2C). The fraction of designs labeling with the FP probe in *E. coli* lysate increased nearly 5-fold from 3% to 17% from round 1 to round 2 (Fig. 2B, Fig. S3). Designs that reacted with the FP probe were purified and incubated with 4MU esters, and two round 1 designs (1.6%) and 10 round 2 designs (5.2%) showed catalytic activity. Retrospective ChemNet analysis of the round 1 designs revealed that the Ser-His H-bonds in the two catalytically active designs were predicted to be among the most preorganized (Fig. S4). ChemNet filtering of round 2 designs on the extent of formation of the key Ser-His H-bond not only increased the fraction of designs exhibiting FP probe labeling and enzymatic activity, but also resulted in higher activities (Fig. 1E,F). The progress curves for these round 1 and 2 designs plateau after approximately one enzyme equivalent of fluorescent product is formed (Fig. 2E), suggesting they catalyze initial nucleophilic attack but fail to hydrolyze the AEI, the rate-limiting step in the cleavage of activated esters^31^. When incubated with substrate, mass spectra of these designs revealed a mass shift corresponding to acylation, further supporting protein inactivation following formation of the acylated intermediate (Fig. S5).

We hypothesized that incorporating a histidine-stabilizing catalytic acid and a second oxyanion hole H-bond donor in a third round of designs (round 3) and filtering for ChemNet preorganization in both the apo and AEI states could generate designs capable of catalytic turnover via hydrolysis of the AEI. For round 3 designs, we required all catalytic triad and oxyanion hole H-bonds to be highly preorganized in ChemNet ensembles of both the apo and AEI states. Of 132 round 3 designs, 111 (84%) displayed FP probe labeling, 20 hydrolyzed 4MU substrates (18%), and two designs (1.5%) displayed multiple turnover activity (Fig. 2B,E). Active designs from all three rounds showed significantly reduced activity upon mutation of any one of the catalytic residues (Ser, His, Asp/Glu, and oxyanion sidechain contact) (Fig. 2E), suggesting that the observed activities are dependent on the designed active site. To determine the kinetic parameters of the active designs, initial or steady-state rates were measured to determine *k*_*2*_/*K*_*m*_ or *k*_*cat*_/*K*_*m*_ for single-turnover and multiple-turnover designs, respectively (Fig. 2E, Fig. S6). For the two designs that displayed catalytic turnover, called ‘super’ and ‘win,’ *k*_*cat*_/*K*_*m*_ values were 22 M^-1^ s^-1^ (*k*_*cat*_ = 0.00137 ± 0.00005 s^-1^, *K*_*m*_ = 64 ± 6 μM) and 410 M^-1^ s^-1^ (*k*_*cat*_ = 0.00117 ± 0.00003 s^-1^, *K*_*m*_ = 2.8 ± 0.3 μM), respectively for the more preferred of the two 4MU substrates (win and super preferentially hydrolyzed 4MU-Ac and 4MU-Bu, respectively (Fig. S7)).

### Structural characterization of designed serine hydrolases

We pursued x-ray crystallography to determine the accuracy with which super and win were designed. We were able to solve crystal structures of both super and win, and found that they had very low Cα RMSDs of 0.8 Å over 165 residues and 0.83 Å over 160 residues (Fig. 3A,D), respectively, to the design models. The very close agreement between experimental and designed structures extends to the geometry of the active site: the sidechain conformations of the catalytic residues are in atomic agreement for super (all-atom RMSD = 0.38 Å over 22 atoms) and for win (all-atom RMSD = 0.86 Å over 20 atoms) except for a rotamer shift in the sidechain oxyanion contact, T99 (Fig. 3B,E). In the active site of super, a water molecule sits above the nucleophilic serine and forms hydrogen bonds with the oxyanion hole contacts, which likely mimics the positioning of the carbonyl oxygen of its ester substrate (Fig. 3B). Similarly, in win, an acetate molecule is positioned at the catalytic center and hydrogen bonds to the Oγ and backbone amide nitrogen of the catalytic serine S142, the oxygen of T99, and the histidine acid/base residue H17, key hydrogen bonds in the catalytic cycle (Fig. 3E).

**Figure 3.**
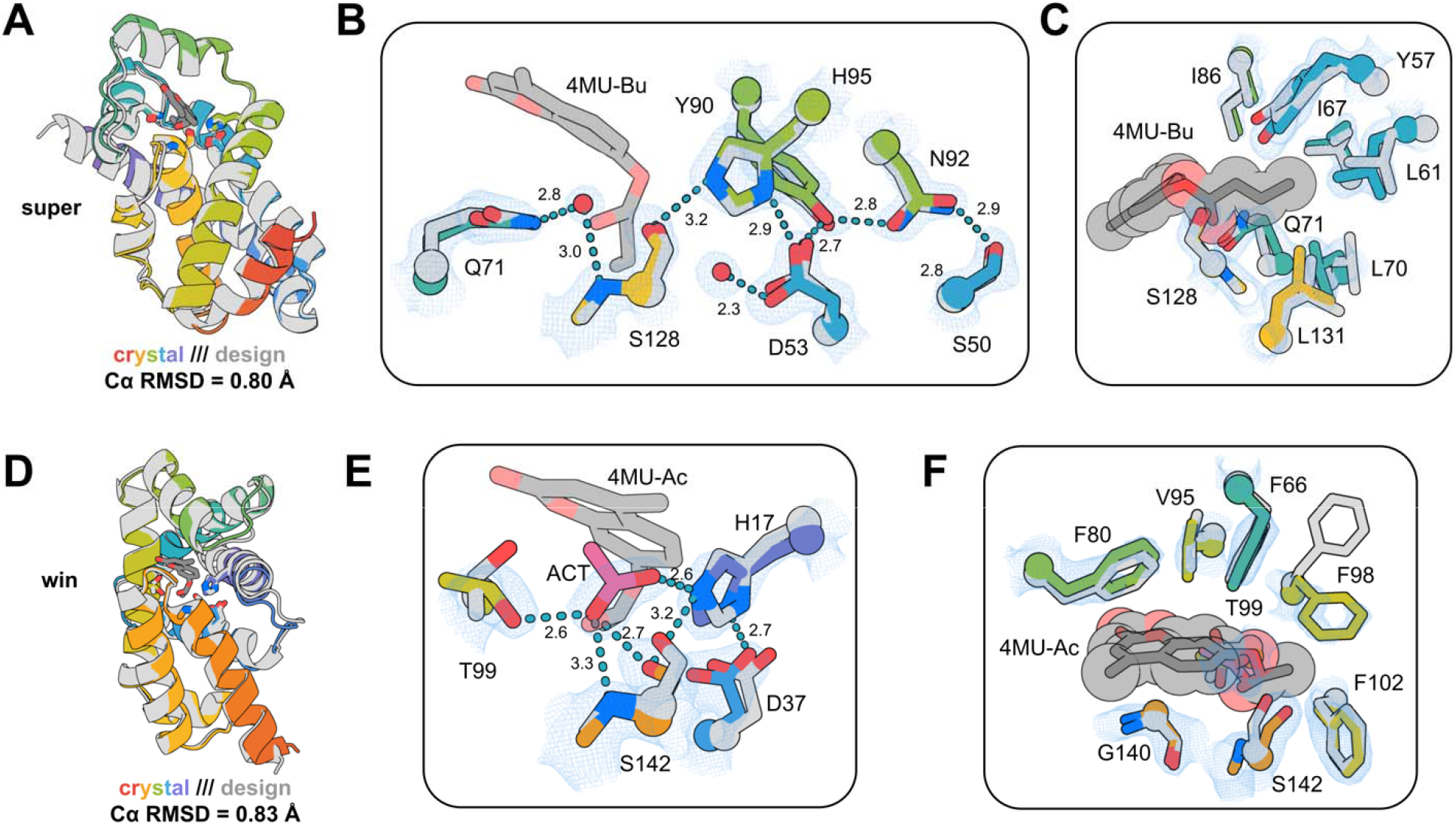
Structural characterization of designed serine hydrolases. **(A**,**D)** Structural superposition of design models (gray) and crystal structures (rainbow) for super (A) and win (D). **(B, E)** Active site overlays of design models (gray) and crystal structures (rainbow) of super (B) and win (E) with 2Fo-Fc map shown at 1σ (blue mesh). **(C, F)** Superposition of substrate binding sites of the design models (gray) and crystal structures (rainbow) with 2Fo-Fc map shown at 1 σ (blue mesh). Distances shown are in Å.

While the structures were solved in the absence of bound small molecule substrate or transition state analogue, overlay of the design model and crystal structure of super reveals high shape complementarity to the butyrate acyl group of its preferred substrate (Fig. 3C and S7). At the same time, the 4MU moiety is largely exposed, corroborating the selectivity of super for 4MU-Bu over 4MU-Ac and suggesting that substrate binding, in this case, is largely driven by binding to the acyl group. For win, a rotamer shift in F98 in the crystal structure would clash with the butyrate moiety, and indeed, win is selective for the smaller substrate 4MU-Ac that avoids this clash (Fig. 3F and S7).

The structures of super and win are very different from known structures; the closest matches found with TM-align to the PDB and larger AlphaFold database have TM-scores of 0.41/0.46 (PDB/AlphaFold database) and 0.46/0.51 (at or below the 0.5 cutoff below which structures are considered to have different topological folds), are proteins of unknown function, and have no similarity to known hydrolases at the fold or active site level (Fig. S8), demonstrating that the design method employed here can find protein structural solutions that extend well beyond those found in nature.

### Filtering for preorganization across the reaction coordinate improves catalysis

We next sought to generate and compare designs filtered explicitly with ChemNet for preorganization over two states (apo and AEI) or over all four states of the reaction path by carrying out additional iterations of LigandMPNN and FastRelax of the active design win (fixing only the identities of the four catalytic residues) (Fig. 4A). We obtained genes encoding 45 two-state filtered designs for experimental characterization, all of which were diverse in sequence compared to the original designs (mean sequence identity to the parent design of 58% and 61% within the active site), and found 38 (84%) labeled with FP-probe. Three of these, win1, win11, and win31, displayed higher *k*_cat_ values compared to the starting design: win has a *k*_cat_ of 0.00117 s^-1^, which increases 15-fold in win1 (0.018 s^-1^), 17-fold in win11 (0.0197 s^-1^), and 9-fold in win31 (0.0105 s^-1^) (Fig. 4B and Fig. S6). Of the 11 four-state filtered designs tested, 10 (91%) labeled with FP-probe (Fig. S9). Two of these, dad_t1 and win_t4, displayed higher catalytic efficiencies than the starting design, with *k*_cat_/*K*_m_ values of 3800 M^-1^ s^-1^ and 640 M^-1^ s^-1^, largely driven by improvements to *k*_*cat*_ (Fig. 4B and S6). Catalytic triad residue knockouts for all designs showed significant reductions in activity. In win11 and win31, mutation of preorganizing residues in the second shell of the active site that H-bond to the catalytic aspartate also significantly reduced activity (Fig. S10).

**Figure 4.**
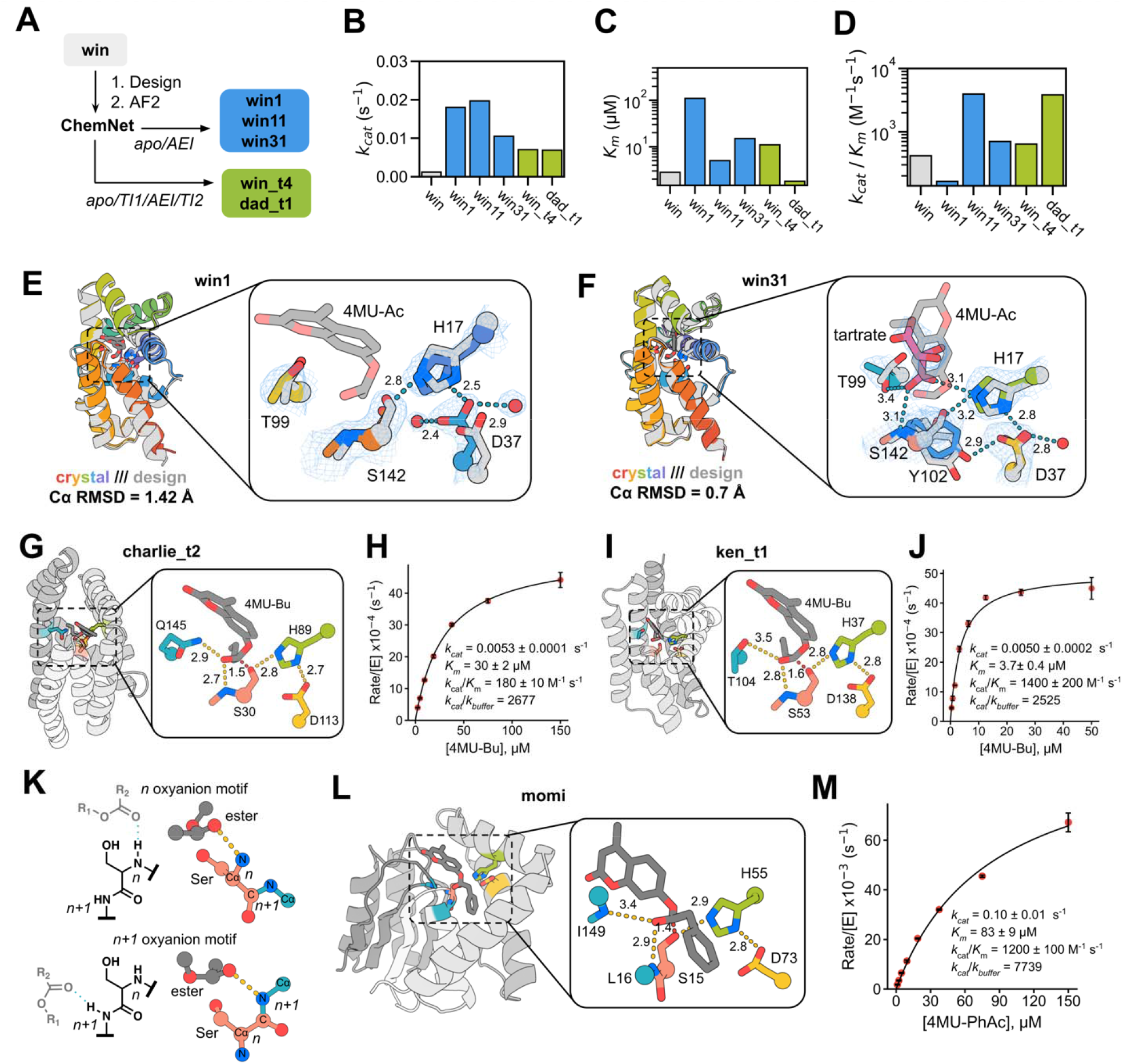
Computational redesign and more complex folds improve catalysis. **(A)** Computational pipeline for redesign of rd3_win. **(B**,**C**,**D)** _*kcat*_ **(B), *K****m***(C)**, and *k*_*cat*_*/K*_*m*_ **(0)** of parent rd3_win compared to computational redesigns. **(E**,**F)** Structural superposition of win1 **(E)** and win31 **(F)** design and crystal structure. **(G**,**H**,**l**,**J)** Design models **(G**,**I)** and Michaelis-Menten plots **(H**,**J)** for active designs with distinct folds and active site structures. **(K)** Chemical and structural comparison of *n* and *n+1* oxyanion hole motifs. **(L)** Design model of active design that utilizes two backbone amide oxyanion hole contacts, one from an n+1 backbone amide. **(M)** Michaelis-Menten plot of active design momi.

We determined the crystal structures of win1 and win31 which revealed very close matches to the design models, with Cα RMSDs of 1.42 Å and 0.7 Å, respectively (Fig. 4E,F). For win1, the active site, including the oxyanion hole sidechain contact, more closely matches the designed conformation (mean all-atom RMSD = 0.54 Å) than the parent design win (Fig. 4E), which may be partly responsible for the 15-fold increase in *k*_*cat*_. For win31, five chains are present in the asymmetric unit, all of which closely match the design model (average Cα RMSD = 0.7 Å) at the backbone level (Fig. 4F and S11). Analysis of the active site across all chains in the asymmetric unit revealed mobility in the catalytic serine, sidechain oxyanion threonine, and a preorganizing tyrosine (Fig. S10), but still a very close match to the design model with a mean all-atom RMSD of 0.7 Å. Tartrate, derived from the crystallization solution, satisfied the electron density present in the active site of all five chains, and forms hydrogen bonds with the serine, histidine, and oxyanion hole contacts (Fig. 4F), likely mimicking key contacts employed throughout the catalytic cycle.

We then explored whether stringent ChemNet filtering for optimal catalytic geometry and preorganization across the reaction coordinate could generate active esterases with novel backbone topologies, active sites, and substrates. We carried out extensive sequence redesign and filtering based on catalytic geometry in all four states starting from round 3 backbones that had not previously displayed esterase activity, and of 20 designs tested, two (charlie_t2 and ken_t1) displayed significant esterase activity, with catalytic efficiencies of 180 M^-1^ s^-1^ and 1400 M^-1^ s^-1^ (Fig. 4G,H,I,J).

To test the generality of this ChemNet filtering approach, we applied it to a different substrate, 4MU-phenylacetate (4MU-PhAc, and a different active site configuration in which the oxyanion hole consists of two backbone amide hydrogen bond donors, rather than a backbone donor and a sidechain donor, and the first backbone donor was the residue following the catalytic serine rather than the catalytic serine itself (Fig. 4K). We used the design pipeline described above to generate 66 designs for this new substrate and catalytic site. The most active of these, momi, displayed a *k*_*cat*_/*K*_m_ of 1240 M^-1^ s^-1^ and *k*_*cat*_ of 0.1 s^-1^, a 5-fold faster rate than win11, the previous best design in terms of turnover number. The distribution of folds generated by RFdiffusion for this active site geometry differed from that for the original geometry, with more α/β fold solutions (as in the case of momi), showing how the RFdiffusion buildup approach crafts overall protein structure topology to the specific active site of interest. The high activity achieved without any prior experimental characterization for this new substrate and catalytic site combination shows that filtering for preorganization throughout the reaction cycle can yield novel catalysts in one shot.

While the catalytic efficiencies of our designed serine hydrolases are far higher than previously reported catalytic triad-based designs, they are still orders of magnitude slower than native hydrolases. Several experimental results identify clear areas for improvement. First, ken_t1 inactivates after roughly 10 turnovers, and mass spectra of the catalyst and the serine knockout incubated with substrate reveal stable acylated species (Fig. S12), indicating that designs that hydrolyze the AEI are still susceptible to inactivation, potentially from off-mechanism acylation events in the active site, which will be important to avoid in future design efforts. Second, in three designs (dad_t1, charlie_t2, ken_t1) from later design rounds made with stringent Chemnet filtering, mutation of the second sidechain oxyanion hole residue has a smaller effect on activity than in the earlier design rounds and compared to analogous mutations in native enzymes (Fig. S10). To investigate the structural effect of the oxyanion hole, we made ChemNet predictions of wild-type and oxyanion hole alanine knockout mutants for all active designs. In the case of super, predictions of Q71A exhibit a clear conformational change of the acylated serine in the AEI which lengthens its distance from the histidine, providing a structural explanation for the loss in activity (Fig. S13). In contrast, wild-type and oxyanion hole knockout predictions were indistinguishable for other designs, including win and high-activity redesigns of win (Fig. S13). Our analysis suggests that the improvements in catalysis achieved throughout our design rounds may derive primarily from improvements in catalytic triad organization and intermediate positioning; future work will focus on optimally placing the oxyanion hole residues to more preferentially stabilize the transition state over the sp^2^ ground state.

### Acyltransferase activity of designed hydrolases

Several native serine hydrolases exhibit promiscuous acyltransferase activity, reacting with small-molecule nucleophiles that compete with hydrolysis to break down the AEI^40^. Due to the long-lived nature of the AEI in these designed hydrolases and the hydrophobicity of their substrate binding pockets, we hypothesized they may also catalyze acyl transfer to aromatic alcohols (Fig. S14). To assess acyl transfer, we incubated designs with their cognate 4MU-ester substrates in the presence of an acyl acceptor, 2-phenylethanol (PEA). For several designs (win, win31, win_t4, and dad_t1), the addition of PEA significantly increased the rate of ester hydrolysis, suggesting these designs catalyzed acyl transfer (Fig. S14). Incubation with PEA and substrate alone or with catalytic serine to alanine knockout mutants of win_t4 and dad_t1 did not exhibit increases in the rate, suggesting observed rate enhancements are enzyme dependent (Fig S14). Acyltransferase activity appears to be anti-correlated with *K*_*m*_: for example, win1 (4MU-Ac K_m_ = 110 μM) was inhibited by PEA, and win (4MU-Ac *K*_m_ = 2.8 μM) had a 3.6-fold maximal rate increase upon addition of PEA, suggesting that transesterification activity may be driven by tighter binding of the acyl acceptor.

### Structural determinants of catalysis

The high structural conservation of catalytic geometry in native serine hydrolases suggests that it is close to optimal for catalysis^32,41^, but it is difficult to assess how activity depends on the detailed geometry of the interactions of the transition states with the catalytic serine, histidine, and oxyanion hole functional groups since while the identities of the catalytic residues can be readily changed by mutation, it is not straightforward to systematically vary backbone geometry. In contrast, our de novo buildup approach samples a wide range of catalytic geometries. To investigate how active site geometry and preorganization influence catalytic activity, we generated ChemNet ensembles of all 812 experimentally characterized designs, categorized as inactive, FP probe labeling, acylation, and catalytic turnover, for each reaction step in the hydrolysis of 4MU-acetate (including design rounds 1-3 and previous NTF2-based designs). The following features were associated with activity.

Increased preorganization and bending of the Ser-His H-bond were associated with higher rates of probe-labeling, acylation, and turnover. All designs capable of catalyzing turnover displayed highly preorganized Ser-His H-bonds across all four states, while inactive designs often displayed rotamer shifts causing loss of the interaction (Fig. 5A,B). Designs that catalyzed turnover had Ser(Oγ):His(Nε-Cε) bond angles that were more acute (median, all states = 94°) than inactive designs (median, all states = 108°), which were more similar to serine-histidine hydrogen bonds across the PDB (∼125°)^33^ (Fig. 5C). This acute H-bond is chemically intuitive given the reaction mechanism, in which this geometry allows histidine to participate, without changing conformation, in all of the necessary proton transfers involving serine, the leaving group oxygen in TI1, and the hydrolytic water^34,42^. This compromise in positioning is observed not only in our active designs but also in many of those found in nature^33,42,43^.

**Figure 5.**
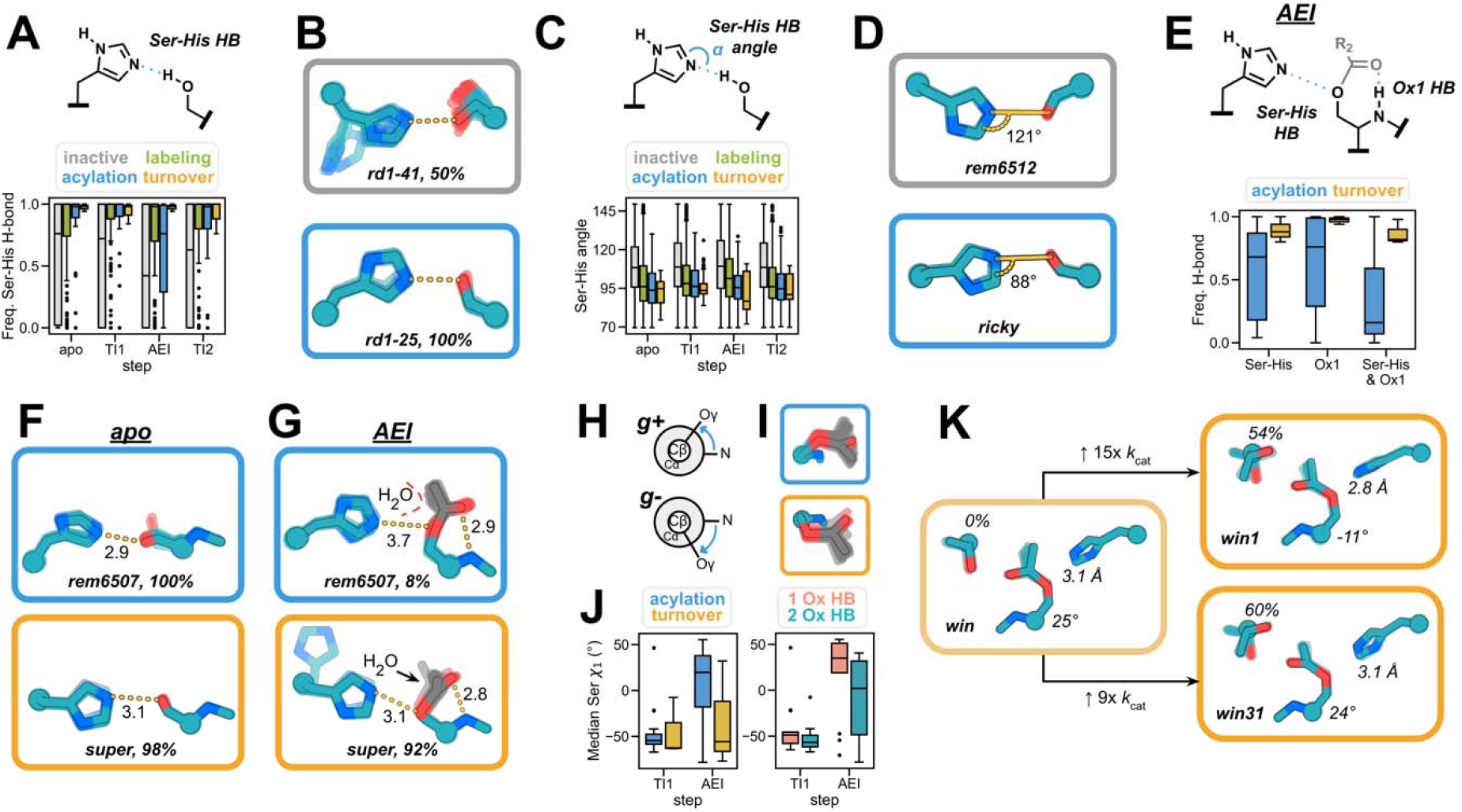
ChemNet ensembles reveal geometric determinants of catalysis. **(A)** Frequencies of catalytic Ser-His H-bond formation in ChemNet ensembles of each reaction intermediate, grouped by experimental outcome. **(B)** Apo ChemNet ensembles of representative inactive (top) and acylating (bottom) designs. **(C)** Median angle (a) between serine Oγ, histidine Nϵ and Cϵ across Chemnet ensembles of inactive and acylating designs. **(D)** Apo ChemNet ensembles of representative inactive (top) and acylating (bottom) designs, angle indicates median *a*. **(E)** AEI ChemNet ensemble H-bond frequencies for designs that undergo acylation or full turnover. **(F)** ChemNet ensembles of the apostate for an acylating (top) and multiple turnover design (bottom). **(G)** ChemNet ensembles of the AEI state for a representative design that undergoes acylation (top) and a design that catalyzes turnover (bottom). Measurements shown represent median distances (A) of key H-bonds indicated for each ensemble and percentages represent frequency of H-bond formation across all ChemNet trajectories. **(H)** Newman projections of serine *g+* and *g-* rotameric states (left). (I) Chemnet ensembles of an acylating design (top) and a design that catalyzes turnover (bottom). **(J)** Median serine *X*_*1*_ angle across Tl1 and AEI state Chemnet ensembles for designs that catalyze acylation or turnover (left). Median serine *X*_*1*_ angle across Tl1 and AEI state Chemnet ensembles for the same designs grouped by number of oxyanion hole hydrogen bonds. **(K)** AEI state Chemnet ensembles for win, win1, and win31, with percent of frames with correct oxyanion hole rotamer, Ser *X*_*1*_ angle, and catalytic Ser-His H-hbond distance shown.

The geometry of the serine rotamer throughout the catalytic cycle was also strongly correlated with experimental outcome. For designs that display acylation or turnover, we found that serine largely occupies the active *g-* rotamer^41^ in the apo state. Designs that display turnover retain the *g-* serine conformer upon formation of the AEI, but designs that irreversibly acylate switch to the *g+* rotamer in the AEI (Fig. 5H,I,J). The *g+* serine rotamer is catalytically incompetent in these designs because it leads to an acyl group conformation that occludes interaction of the hydrolytic water with histidine (Fig. 5G), increases the median Ser-His H-bond distance (Fig. 5G), and reduces the frequency that the Ser-His and oxyanion hole-acyl group H-bonds form (Fig. 5E). The same retention of the *g-* rotamer in the AEI is observed in native crystal structures^34^. ChemNet analysis also revealed that the presence of a second oxyanion hole residue favors the active *g-* serine rotamer: those designs with only one oxyanion hole H-bond (from the backbone amide of the serine nucleophile) shift from *g-* to *g+* upon acylation, while designs with two oxyanion hole H-bonds predominantly occupy *g-* Ser rotamers (Fig. 5J, right). The second oxyanion hole contact in serine hydrolases thus not only stabilizes the transition state but likely helps orient intermediates in catalytically productive conformations.

Differential preorganization may also explain activity trends in the win, win1, and win31 series. ChemNet analysis of the crystal structures of these designs revealed that in the AEI state, the more active win1 and win31 sample the designed T99 oxyanion hole rotamer in 56 and 60% of predictions, respectively, while win never adopts this rotamer (Fig. 5K). Although both observed rotamers place T99 Oγ within hydrogen bonding distance of the oxyanion, the designed rotamer-oxyanion dihedral angle (91°) much more closely matches the angles observed in native serine hydrolases, suggesting it is likely more optimal for selective transition state stabilization^33,44,45^ (see Methods). We also observed differences in the serine rotameric state and the preorganization of the acyl group in the AEI state. Both win and win31 occupy the catalytically unfavorable *g+* rotamer across the entire AEI ensemble, while win1 displays a less pronounced rotameric shift, which leads to shorter serine-histidine hydrogen bond distances (2.8 Å in win1 compared to 3.1 Å in win and win31). Overall, the acyl groups of win1 and especially win31 display significantly less conformational heterogeneity than that of win, which presumably increases the likelihood of histidine-mediated water attack (Fig. 5K).

## Conclusions

The substantial catalytic efficiencies of 10^3^ M^-1^ s^-1^, the complexity of the active site geometry, and the accuracy of sidechain placement considerably surpass previous serine hydrolase computational design efforts despite the testing of a relatively small number of designs and complete omission of laboratory optimization. The folds of the designed catalysts are very different from those of natural serine hydrolases, demonstrating the ability of generative deep learning design methods to find completely new solutions to design challenges that differ from those found by natural evolution. Previous efforts to design catalytic triad-based designs have failed to achieve multiple turnover; in some cases, such as our preliminary NTF2-based designs, a backbone amide oxyanion hole was impossible to achieve due to scaffold limitations, while in others based on native scaffolds, the histidine geometry was difficult to control which limited activation of the leaving groups and water (Fig. S15)^8^. De novo backbone generation building outward from a specified active site with RFdiffusion overcomes these limitations by enabling generation of almost any desired catalytic geometry.

Assessing design compatibility over the entire catalytic cycle has been a longstanding challenge in enzyme design. We show that the deep neural network ChemNet can rapidly generate ensembles for a series of reaction intermediates which directly assess preorganization, and provide structural insights that would otherwise require labor-intensive structural studies. For example, ChemNet revealed pervasive off-target conformational changes in the acyl-enzyme, which could be responsible for the failure to catalyze turnover for many previously designed esterases. The stochastic nature of ChemNet provides ensemble views of the energy landscapes around key reaction intermediates; the agreement we observe between ChemNet preorganization and experimental success rates suggests that such ensemble generation will be broadly useful for enzyme design moving forward.

While the designed catalysts described here are far more active than previous de novo designed serine hydrolases obtained by direct computation, they are still two to three orders of magnitude less efficient than native serine hydrolases, particularly in terms of turnover number. There are several possible explanations for the remaining activity gap: (1) the oxyanion hole identities and geometries differ slightly from those in native structures, which could reduce selective transition state stabilization,^33,44,45^ (2) the catalytic aspartate in the designs rarely participates in 2-3 additional hydrogen bonds (like those found in nature) which may limit its modulation of the catalytic histidine’s pK_a_, and (3) the designed active sites are more buried than those of natural serine proteases, which could inhibit water entry into the active site for acylenzyme hydrolysis. Our de novo buildup approach using RFdiffusion coupled with ChemNet ensemble analysis to ensure design accuracy and preorganization should allow us to test all of these hypotheses by direct construction, which should further complement more traditional approaches based on structural examination and mutation of highly evolved native enzymes.

More generally, we anticipate that the ability to precisely position multiple catalytic groups with sub-angstrom precision using RFdiffusion, and to assess active site organization throughout a complex reaction cycle using ChemNet should enable the design of a wide variety of new catalysts in the near future.

## Acknowledgments

We thank Luki Goldschmidt and Kandise VanWormer for maintaining the computational and wet lab resources at the Institute for Protein Design. We thank Anthony P. Green, Florence J. Hardy, and Donald Hilvert for helpful advice during the development of this project. We thank Florence J. Hardy and Madison A. Kennedy for reading and editing drafts of the manuscript.

## Author contributions

Conceived the study: A.L., S.J.P., and D.B; Trained chemnet: I.A. Conceived, implemented, and trained the models comprising all-atom CA RFdiffusion described here: D.J.; Implemented code to support training of all-atom RFdiffusion models: W.A.; Performed DFT calculations used to model the substrate geometry for design calculations: C.J.; Developed motif generation script: I.K.; Computationally designed serine hydrolases: A.L., S.J.P, A.S., A.H., and K.H.S.; Experimentally characterized serine hydrolases: A.L., S.J.P, K.H.S., A.S., A.H.; Prepared samples for crystallography: A.L. and S.J.P.; Performed crystallization and crystal preparation: E.B. and A.K. Performed data collection for crystal structures: A.B. and B.S.; Solved and refined crystal structures: A.L., S.J.P, and A.B. Wrote the manuscript: A.L., S.J.P, I.A., and D.B.; All authors revised and edited the manuscript. Supervision: D.B. and K.H.

## Methods

### Computational design of serine hydrolases

#### Motif generation

Motifs were built in an iterative process. First, a substrate rotamer in a transition state geometry (either 4MU-Bu or 4MU-Ac) was placed in accordance with geometries in ref 7 in relation to a 3-residue stub of the serine and local oxyanion hole from one of two natural serine hydrolase crystal structures (1scn, residues 220-222, and 1lns, residues 347-349, in which all residues other than the serine were mutated to alanine). The transition state geometry of the substrate ester group was determined by DFT geometry optimization (B3LYP-D3(BJ)/6-31G(d)). Next, positions and rotamers of histidine on 3-residue helical or strand stubs flanked by alanine were sampled around the catalytic serine and filtered for those structures in which the histidine simultaneously formed hydrogen bonds with the catalytic serine and the substrate leaving group oxygen. This process resulted in 108 unique round 1 motifs. For the round 3 motifs, initially the aspartate or glutamate residue and second oxyanion hole hydrogen bond were added in a similar manner using geometric sampling of hydrogen-bonding conformations and rotamers. However, backbones produced from these motifs had exceedingly low AF2 success rates, presumably due to the generation of incompatible combinations of backbone conformations. To ensure that the remaining catalytic residue stubs were placed in realizable geometries, we generated 10,000 backbones with RFdiffusion using the simple substrate-Ser-His motifs as input, and then searched these backbones using Rosetta for positions on secondary structure that could accommodate the aspartate or glutamate triad residue to hydrogen bond to histidine. These stubs were then extracted, and in a final step, the same process was repeated to generate stubs for the second oxyanion hole, considering all hydrogen bond donating sidechains, ultimately producing 2238 unique round 3 motifs with Ser-His-Asp/Glu catalytic triads, and Ser/Thr/Tyr/His/Trp oxyanion holes.

#### Backbone generation

See supplemental methods for a detailed description of CA diffusion, which was employed to generate backbones to scaffold motifs.

#### Sequence design

We performed three cycles of LigandMPNN^36^ and Rosetta FastRelax^46^ to design sequences for backbones generated from RFdiffusion. To encourage formation of hydrogen bond contacts to the catalytic histidine (for round 1 motifs) and to the catalytic aspartate/glutamate (round 3 motifs), the log probabilities used by LigandMPNN to select residues were biased toward polar amino acids for all residues with Cα within 8 Å of the active site. Catalytic residues were kept fixed and Rosetta enzyme constraints^47,48^ were applied during the relax steps to maintain the catalytic geometry during cycles of design. Constraints were defined for each hydrogen bonding interaction using the starting motif geometry with tolerances of 0.1 Å for distances and 5° for angles and dihedrals.

#### Filtering

After sequence design, designs were filtered on the recapitulation of the motif catalytic geometry after FastRelax and the shape complementarity of the binding site to the substrate. Sequences of passing designs were used as input to AF2^39^ for single sequence structure prediction. AF2 was run using model 4 with three recycles. Designs were filtered for a global Cα RMSD < 1.5 Å, pLDDT > 75, and catalytic residue Cα RMSD < 1.0 Å. Designs that passed AF2 filters were subsequently analyzed using ChemNet. ChemNet is a denoising neural network which was trained on high- and medium-resolution X-ray and EM structures from the PDB to recapitulate the correct atom positions from partially corrupted input structures provided that all the chemical information about the system being modeled is known from the start. ChemNet predictions were done for a spatial crop of 600 atoms closest to the active site. The inputs to the network included the protein backbone coordinates within the crop and the amino acid sequence with side chain coordinates randomly initialized around the respective C-alpha atoms. For proteins without a crystal structure, the AF2 model was used. For every designed protein, we modeled 5 reaction states differing in the chemical modifications the catalytic serine undergoes in the course of the reaction: (1) apo, (2) substrate bound, (3) tetrahedral intermediate 1, (4) acylenzyme intermediate, and (5) tetrahedral intermediate 2. We used 50 different seeds to generate an ensemble of 50 ChemNet models for each reaction state (apo, substrate bound, TI1, AEI, and TI2). These ensembles were then individually analyzed for the preservation of hydrogen bonding patterns in the active site. For each of the 50 predictions in each ensemble, geometries of each hydrogen bonding interaction in that step (see Supplemental Methods) were measured. To analyze native hydrolases with Chemnet, a set of native crystal structures was collected^33^ (PDB IDs: 1ACB_E, 1C5L_H, 1H2W_A, 1IC6_A, 1IVY_A, 1PFQ_A, 1QNJ_A, 1QTR_A, 1ST2_A, 2H5C_A, 2QAA_A, 3MI4_A, 5JXG_A), the active site locations identified, and the above-described process was applied.

### In-gel fluorescence screening with activity-based probes

DNA encoding the designed proteins was ordered from IDT as eblocks and cloned into vector LM627 (addgene), which contains a C-terminal SNAC and hexahistidine tag. Resulting plasmid was transformed into BL21(DE3) cells and grown overnight in 1 mL of LB supplemented with 50 μg/ml kanamycin. For expression, 100 μL of overnight was used to inoculate 1 mL of LB media and grown for 1.5 hours at 37°C on a Heidolph shaker and then 10 μL of 100 mM IPTG was added and cultures were shaken at 37°C for an additional 3 hours. Cultures were centrifuged at 4000*g* for 10 minutes and supernatant removed. Cell pellets were resuspended in 200 µL 20 mM HEPES (pH 7.4), containing 50 mM NaCl, 0.1 mg/mL lysozyme, and 0.01 mg/mL DNaseI. After 15 minutes, lysates were frozen in liquid nitrogen and subsequently thawed. 10 µL of lysate was incubated with 1 µM FP-TAMRA probe (10 µL of 2 µM stock in lysis buffer) for 1 hour at room temperature before quenching using 2x Laemmli sample buffer. Labeled samples were heated at 95°C for 5 minutes and 10 µL of each sample was separated on a BioRad AnykD Criterion precast gel and in-gel fluorescence was visualized using a LI-COR Odyssey M imager. Gels were subsequently stained with coomassie blue to visualize the molecular weights and levels of expression of each design.

### Lysate screening

DNA encoding the designed proteins was ordered from IDT as eblocks and cloned into vector pCOOL1 which contains a C-terminal mScarlet-i3 fusion and His tag. Cultures were grown overnight at 1 mL scale in 96-well plates on a Heidolph shaker at 1300 rpm and 37 °C. For expression, 50 μL of the overnight cultures were used to inoculate 1 mL of autoinduction media in 96-well round bottom plates and incubated at 1300 rpm and 37 °C for approximately 24 hours. Cultures were centrifuged at 4000*g* for 10 minutes and supernatant decanted, followed by a wash with buffer (20 mM HEPES, 50 mM NaCl, pH 7.4) and incubation on a Heidolph shaker at 1300 rpm at room temp for 5 minutes to resuspend. Plates were centrifuged again at 4000*g* for 10 minutes and supernatant decanted. For lysis, cell pellets were resuspended with 500 μL of lysis buffer (20 mM HEPES, 50 mM NaCl, 0.01 mg/mL DNAseI, 0.01 mg/mL lysozyme, 1 mM EDTA, 0.1% triton X-100) and incubated for 2 hours on a Heidolph shaker at 1300 rpm and 37 °C. Plates were centrifuged at 4300*g* for 30 minutes and supernatant collected for screening. For activity screening, 4 or 6 μL of lysate was aliquoted into microtiter plates and reactions initiated by addition of 36 or 54 μL of buffer containing 111.1 μM 4MU-Ac or 4MU-Bu, 20 mM HEPES, 50 mM NaCl, pH 7.4, 5% DMSO.

### Protein expression and purification

Genes encoding the designed proteins were ordered from IDT as eblocks and cloned into vector LM627 (addgene) (ref). Resulting plasmid was transformed into BL21(DE3) cells and grown overnight in 1 mL of LB supplemented with 50 μg/ml kanamycin, after which 500 μL of overnight was used to inoculate 50 mL of autoinduction media, which was grown 4-6 hours at 37 °C and then overnight at 18 °C. Cultures were spun down at 4000*g* for 15 minutes, and supernatant decanted. Cell pellets were resuspended in 25 mL of cold wash buffer (40 mM imidazole, 500 mM NaCl, 50 mM sodium phosphate, pH 7.4) with 1 mg/mL lysozyme and 0.1 mg/mL DNAse I. Cell slurries were sonicated on ice for 2.5 minutes at 80% amplitude, 10s on 10s off. The resulting lysate was centrifuged at 14000*g* for 30 minutes and the supernatant was applied to 1 mL of Ni-NTA resin equilibrated with wash buffer. The resin was subsequently washed with 15 mL of wash buffer 3 times and once with 400 μL of elution buffer (400 mM imidazole, 500 mM NaCl, 50 mM sodium phosphate, pH 7.4) followed by elution with 1.3 mL elution buffer. The eluate was purified by size-exclusion chromatography on a Superdex 75 Increase 10/300 GL with running buffer of 20 mM HEPES, 50 mM NaCl, pH 7.4. Samples were either used immediately in downstream experiments or snap frozen in liquid nitrogen and stored at -80 C. Protein molecular weight was confirmed by LC-MS.

### Kinetic analysis

To characterize hits identified from in-gel fluorescence and lysate screens for catalytic turnover, we incubated purified protein samples with fluorogenic substrates 4MU-Ac, 4MU-Bu and 4MU-PhAc. Kinetic screens were either performed in 40 μL reaction volumes in 96-well half area plates or 60 μL reaction volume in 96-well full-area plates. Protein and substrate were prepared in 20 mM HEPES, 50 mM NaCl, pH 7.4, 5% DMSO. Either 4 or 6 μL of enzyme was added to microtiter plates and the reactions were initiated by addition of substrate (36 or 54 μL). Generation of the fluorogenic product 4MU was monitored continuously (excitation 365 nm, emission 445 nm). Analysis of the resulting data were carried out using custom scripts (see computational methods). In cases where single-turnover activity was observed, initial velocities were used to determine *k*_2_/*K*_m_. For those designs that displayed a clear burst phase followed by a slower steady-state rate, straight-line fits of the steady-state velocities were used to determine Michaelis-Menten catalytic parameters.

To determine the uncatalyzed reaction rate in assay buffer (20 mM HEPES, 50 mM NaCl, pH 7.4, 5% DMSO), substrate was diluted in buffer alone and rates determined at multiple substrate concentrations, after which the rate was determined from fitting [S] versus rate with an equation of the form rate = *k*_buffer_[S].

### Crystallography

Proteins for crystallography were prepared as described above, but SEC was done with SNAC tag cleavage buffer^49^. After SEC, protein eluate was incubated with 500 mM guanidinium hydrochloride and 2 mM NiCl_2_ overnight at room temperature to remove the C-terminal His tag. The SNAC cleavage reaction was applied to a nickel column equilibrated with wash buffer to remove any uncleaved product and resulting eluate applied to a Superdex 75 Increase 10/300 GL column with 20 mM HEPES, 50 mM NaCl, pH 7.4 as the running buffer. Samples were concentrated and stored at -80° C or immediately used for crystallization. Crystallization screening was performed using a Mosquito LCP by STP Labtech and resulting crystals were harvested directly from the screening plate. Crystallization conditions for each design were as follows: slap215.8 (15 mg/mL) in 0.1 M Bis-Tris pH 5.5, 25% (w/v) PEG 3350, super (50 mg/mL) in 0.2 M Potassium fluoride, 20% (w/v) PEG 3350, win (42 mg/mL) in 0.1 M Sodium acetate pH 4.6, 8% (w/v) PEG 4000, win1 (54 mg/mL) in 60% v/v Tacsimate pH 7.0, and win31 (60 mg/mL) in 0.2 M di-Ammonium tartrate and 20% (w/v) PEG 3350. Data were processed with XDS^50^, phased and refined with Phenix^51^, and model building performed with COOT^52^. Coordinates are deposited in the PDB with PDB IDs of 9DED (slap215.8), 9DEE (super), 9DEF (win), 9DEG (win1), and 9DEH (win31).

### Mass spectrometry

Intact mass spectra of protein samples were obtained by reverse-phase LC/MS on an Agilent G6230B TOF after desalting using an AdvanceBio RP-Desalting column. Deconvolution using a total entropy algorithm was performed using Bioconfirm. In some cases, protein samples (1 mg/mL) were incubated overnight with substrate (300 μM) in SEC running buffer at room temperature prior to mass spectrometry analysis.

### Acyltransferase activity screening

Enzymes (1 μM) were incubated with 100 μM cognate substrate in assay buffer in the presence of varying concentrations of acyl acceptor, PEA (50, 25, 12.5, 6.3, 3.1, 1.6, 0.8, 0 mM), and substrate hydrolysis were monitored for 1 hour as described above. Initial velocities were obtained by fitting the beginning of each progress curve and divided by the hydrolysis rate in the absence of PEA to obtain relative rates of hydrolysis.

### Structural similarity search of the PDB and AFDB

To assess the structural novelty of our designed enzymes, we used TMalign^53^ to compare our crystal structures against the Protein DataBank (PDB) and AlphaFold database^54^. We downloaded all protein polymers from the PDB solved by X-ray crystallography or Cryo-EM on April 4, 2024 and extracted all protein chains from each entry. Models of AFDB50 ^55^ (version 4) proteins were fetched April, 2024. We report the average TM-score for the top hit.

