## Supplemental Materials for "Computational design of serine hydrolases"

**Table 1. Data collection and refinement statistics.**

|  | **slap215.8** | **super** | **win** | **win1** | **win31** |
| --- | --- | --- | --- | --- | --- |
| **Wavelength** | 0.97648 | 0.92010 | 1.00002 | 1.00002 | 0.97936 |
| **Resolution range** | 40.00 - 1.77 (1.81 - 1.77) | 32.12 - 1.21 (1.28 - 1.21) | 46.02 - 1.75 (1.78 - 1.75) | 43.27 - 2.11 (2.17 - 2.11) | 58.33 - 2.21 (2.33 - 2.21) |
| **Space group** | P 1 21 1 | R 3 :H | P 21 2 21 | P 21 21 21 | P 21 21 21 |
| **Unit cell** | 38.55, 31.55, 40.79; 90, 101.32, 90 | 68.42, 68.42, 76.44; 90, 90, 120 | 31.16, 46.02, 92.94; 90, 90, 90 | 35.81, 82.03, 101.88; 90, 90, 90 | 63.40, 106.99, 116.66; 90, 90, 90 |
| **Total reflections** | 63121 (3594) | 2592198 (38409) | 153901 (8688) | 192423 (15480) | 457352 (66573) |
| **Unique reflections** | 9440 (526) | 40645 (5957) | 14105 (749) | 17989 (1414) | 40598 (5849) |
| **Multiplicity** | 6.7 (6.8) | 6.4 (6.4) | 10.9 (11.6) | 10.7 (10.9) | 11.3 (11.4) |
| **Completeness (%)** | 98.8 (99.3) | 99.86 (100.00) | 99.80 (99.60) | 99.9 (99.9) | 100.00 (100.00) |
| **Mean I/sigma(I)** | 11.4 (2.4) | 12.6 (1.1) | 14.0 (1.9) | 10.3 (2.4) | 6.4 (0.70) |
| **Wilson B-factor** | 22 | 21 | 27 | 28 | 39 |
| **R-merge** | 0.090 (0.831) | 0.048(1.347) | 0.084 (1.274) | 0.151 (0.958) | 0.284 (3.183) |
| **R-pim** | 0.041 (0.368) | 0.022 (0.639) | 0.028 (0.402) | 0.050 (0.314) | 0.090 (1.024) |
| **CC1/2** | 0.997 (0.798) | 0.999 (0.471) | 0.998 (0.752) | 0.999 (0.918) | 0.977 (0.424) |
| **Reflections used in refinement** | 9422 (1332) | 40645 (2937) | 14057 (1366) | 17872 (1321) | 40457 (2855) |
| **Reflections used for R-free** | 935 (138) | 2012 (150) | 1406 (137) | 1788 (131) | 2004 (138) |
| **R-work** | 0.1896 (0.2479) | 0.2057 (0.3380) | 0.2361 (0.3217) | 0.2039 (0.2557) | 0.2381 (0.3796) |
| **R-free** | 0.2216 (0.2904) | 0.2414 (0.3917) | 0.2726 (0.3556) | 0.2590 (0.3229) | 0.2697 (0.3863) |
| **Number of non-hydrogen atoms** | 1044 | 1451 | 1284 | 2700 | 6346 |
| **macromolecules** | 969 | 1292 | 1215 | 2535 | 6204 |
| **ligands** | 0 | 0 | 5 | 6 | 83 |
| **solvent** | 75 | 159 | 64 | 159 | 59 |
| **Protein residues** | 114 | 160 | 151 | 314 | 790 |
| **RMS(bonds)** | 0.019 | 0.006 | 0.007 | 0.011 | 0.008 |
| **RMS(angles)** | 1.62 | 0.60 | 0.81 | 1.03 | 0.75 |
| **Ramachandran favored (%)** | 98.21 | 99.37 | 97.99 | 99.68 | 99.36 |
| **Ramachandran allowed (%)** | 1.79 | 0.63 | 2.01 | 0.32 | 0.64 |
| **Ramachandran outliers (%)** | 0.00 | 0.00 | 0.00 | 0.00 | 0.00 |
| **Average B-factor** | 26 | 27 | 40 | 33 | 49 |
| **macromolecules** | 26 | 26 | 40 | 33 | 49 |
| **ligands** |  |  | 34 | 51 | 56 |
| **solvent** | 315 | 35 | 35 | 36 | 40 |

Statistics for the highest-resolution shell are shown in parentheses

**Motif generation via conformational sampling**

To generate constellations of histidine rotamers around the serine backbone motif, distances, angles, and torsions were first specified in enzyme constraint file format along with sampling ranges and frequencies for the serine-substrate and serine-histidine interactions. These were input to a script which (1) sampled probable histidine rotamers from the Dunbrack library at the specified geometries with respect to the serine nucleophile and substrate, (2) built out protein stubs in helical or strand conformation flanking the histidine using Rosetta, (3) and filtered for clashes between the substrate, serine stub, and histidine stub before outputting these conformations as PDB files. The code and a detailed description of this script can be found here: <https://github.com/ikalvet/invrotzyme> .

**Evaluation of hydrogen bonds in ChemNet predictions**

Hydrogen-bonding interactions in ChemNet predictions were evaluated by the measuring of distances, angles, and torsions between the acceptor and donor heavy atoms. Upper and lower bound cutoffs for each parameter (distance, angle, dihedral) were used to determine if the interaction constituted a hydrogen bond, depending on the hybridization of the participating heavy atoms.

**Structure generation with a new variant of all-atom RFdiffusion.**

In this study, we follow the familiar protocol of generating protein backbone structures with a diffusion model, followed by a fixed backbone sequence design step. The diffusion model we use is a newly trained variant of RFdiffusionAA, which we call ‘CA RFdiffusion’ (as in, alpha-carbon). In this section, we detail the training and inference protocols for this model. All training and inference code will be released upon manuscript publication.

**Introduction**

CA RFdiffusion comprises a set of two models, a diffusion model and a “refinement” model, that are both fine-tuned versions of the same pretrained RosettaFold All-Atom structure prediction network. There are two main differences between CA RFdiffusion and any previous variant of RFdiffusion(AA). First, the diffusion model itself produces only alpha-carbon coordinate positions, instead of full N-CA-C frames with both a translation and orientation (as in RFdiffusion[[1]](https://paperpile.com/c/IXs5yq/m5gK) and RFdiffusionAA[[2]](https://paperpile.com/c/IXs5yq/Wd6r)). This means that a second step after generating the “CA trace” must be taken to obtain a fully constrained polypeptide backbone which can be assigned a sequence (one could also design sequences from CA coordinates alone, but we did not attempt this). We perform a second “refinement” step by training a second separate model to produce a full protein backbone given just the trace of alpha carbons.

The second main difference is that instead of taking in motif structure information via static 3D coordinates supplied to the network (as is the case in RFdiffusion and RFdiffusionAA), CA RFdiffusion takes in motif structure information via the available inter-residue pairwise distance and orientation input. In RosettaFold All-Atom (and other previous structure prediction networks) this input is normally used for supplying structural information of proteins homologous to the query sequence during structure prediction (see [[2], [3], [4]](https://paperpile.com/c/IXs5yq/Wd6r+drdi+dSMX)). For CA RFdiffusion, we use this input to encode motif structures as their pairwise distances and orientations, and have the network reconstruct the motif in its output 3D predictions. It is noteworthy that at least one other group [[5]](https://paperpile.com/c/IXs5yq/dQPa) has developed an analogous motif-scaffolding technique concurrently and independently of the method we describe here. One main benefit of this motif input style is that, as Lin et al. [[5]](https://paperpile.com/c/IXs5yq/dQPa) also point out, a user is able to present multiple discontiguous motifs to the network for scaffolding without specifying their relative rigid body transform (as is mandatory, by definition, for methods that take in 3D coordinates of the motif).

**Training CA RFdiffusion**

Here we detail the two-part training of CA RFdiffusion, which comprises training a diffusion model for generating CA traces, followed by training a “refinement” model which produces full protein backbones from the CA traces.

**Training the diffusion model**

**Architecture**

Training the diffusion model for CA RFdiffusion is similar to [[1]](https://paperpile.com/c/IXs5yq/m5gK), but there are several differences in the inputs to the network and loss function.

We begin with the architecture and pretrained weights of RoseTTAFold All-Atom [[2]](https://paperpile.com/c/IXs5yq/Wd6r). To this architecture, we expand the dimensions of the template inputs (see Krishna et al. 2024 for architecture details) to include one additional per-token and per-token-pair indicator feature. These denote whether or not a token, or pair of tokens, is part of a motif being scaffolded by the network. During training, we supply three distinct templates (analogous to three separate homologous structure inputs during structure prediction), as shown in Table 2.

|  | **Per-token (1D) info** | **Per-token-pair (2D) info** |
| --- | --- | --- |
| **Template #1 -** Self-conditioning | **Timestep feature:** 1-T/t  **Is motif feature:** -1 | **Structural data:** Previous prediction of X_0_  Pair constraint indicator: -1 |
| **Template #2 -** Current X_t_ input | **Timestep feature:** -1  **Is motif feature:** -1 | **Structural data:** Current X_t_ structure input  Pair constraint indicator: -1 |
| **Template #3 -** Perfect motif info | **Timestep feature:** -1  **Is motif feature:** 0 for False, 1 for True | **Structural data:** Perfect motif geometry  **Pair constraint indicator:** 1 if token pair is geometrically constrained, else 0. |

***Table 2:*** *Template indicator, timestep and structural features input to both the diffusion and refinement models of CA RFdiffusion. Template #1 is the self conditioning template, exactly as described in* [*[1]*](https://paperpile.com/c/IXs5yq/m5gK)*. For the new per token “is motif” indicator, this template receives a null value of -1 for all tokens, while still containing the encoding of the current discrete time step 1-T/t. The pairwise template features for template #1 contain the self-conditioning information (the distance/angle encoding of the previous prediction of X_0_), and a null -1 indicator feature for all pairs. Template #2 contains null information (i.e., -1) for all indicator and timestep features, but the pairwise structure input contains the distance/angle encoding of the current noisy X_t_ structure being input to the network in 3D. Template #3 contains information about the perfect motif geometry and primary sequence location, to be scaffolded by the network. This template is the only one of the three which uses the “is motif” feature (0 if non-motif, 1 if motif token) and the “pair is constrained” indicator (1 if the pair is constrained, else 0).*

We initialize the weights associated with the expanded feature dimensions as all zeros, such that predictions with the expanded architecture completely ignore the new features at first. However, as training progresses, the network can learn to correlate the motif indicator features with the perfect motif information supplied in template #3, helping distinguish that particular structural input from the rest of the structural inputs.

**Preparing training examples**

Training examples are prepared in two stages: First, a dataset is selected to draw a structure from; either (A) PDB protein monomers without small molecules, or (B) PDB protein monomers in complex with small molecules. Then, a masking strategy is chosen for the drawn structure. The masking strategies available to either dataset are not the same, and they are outlined in Table 3 below.

| **Dataset probabilities** | **Masking probabilities** |
| --- | --- |
| Dataset #1: PDB monomers - 60% probability | (A) “Chunked diffusion mask” - 20%  (B) Multi triple contact - 50%  (C) Unconditional - 30% |
| Dataset #2: PDB monomers in complex with ligands - 40% probability | (A) Small molecule contact mask - 100% |

***Table 3:*** *Training dataset and masking strategy probabilities for diffusion model training. For protein-only monomers from the PDB, which is chosen 60% of the time, there are three possible masking strategies. (A) A “chunked” diffusion mask, in which 1-8 discontiguous fragments are designated as motif components. Each pair of discontiguous fragments has some nonzero probability of being unconstrained with respect to each other (see Algorithm 1 for details). (B) Multiple “triple contacts” are chosen, in which three residues that are close in spatial proximity but far from each other in primary sequence are selected (see Algorithm 2 for details). (C) Unconditional, in which there are no motifs and the task is to denoise the full structure. For the second dataset, which is PDB monomers in complex with small molecule ligands, there is a single motif masking strategy used every time. In this “small molecule contact mask” generation, the structure of the ligand is revealed in the template, along with either 0, 1, or 2 additional discontiguous protein fragments in close proximity to the ligand (see Algorithm 3 for details).*

Once the dataset and masking strategy are chosen, any structural components which are considered motif (any ligand, and unmasked protein fragments) are encoded in the motif (third) template with appropriate indicator features.

To noise a selected example, we uniformly sample from the discrete set of times t ∈ [1,200] and then adopt precisely the CA-coordinate noising strategy employed by [[1]](https://paperpile.com/c/IXs5yq/m5gK) to noise the CA positions of each N-CA-C frame in the backbone and all atoms in any ligand. We do not use any noising strategy for the frame orientations, and instead set all residue frame orientations to the identity rotation, which removes any information about the native structure contained in the frame orientations. It is noteworthy that this step breaks the equivariance of a network forward pass, because the process of setting all frame orientations to an arbitrary orientation is not equivariant to arbitrary rotations of the input molecules. A summary of training example generation hyperparameters can be found in Table 4. It is worth emphasizing that because the motif information is encoded in the template inputs to the network (described above), all residues and small molecule atoms are noised in 3D space, and the network is tasked with reconstructing the motif in its output--a well defined problem since the motif is templated.

| **Parameter** | **Description** | **Value** |
| --- | --- | --- |
| β_0_ | Variance at time t=0 | 0.01 |
| β_T_ | Variance at time t=T=200 | 0.07 |
| T | Total number of timesteps in noising process | 200 |
| Variance schedule type | NA | Linear |
| W_disp_ | CA displacement loss weight | 0.5 |
| W_dist-ang_ | Distogram and anglogram cross entropy loss weight | 0.05 |
| W_FAPE,prot_ | Intra-protein fragment FAPE loss weight | 10.0 |
| W_FAPE,prot-sm_ | Inter protein-ligand FAPE loss weight | 10.0 |
| W_FAPE,sm_ | Intra ligand FAPE loss weight | 10.0 |
| W_FAPE,prot(non-motif)_ | (Refinement model only) Non-motif protein FAPE loss weight | 10.0 |
| W_pLDDT_ | (Refinement model only) pLDDT loss weight | 0.1 |
| A_FAPE,prot_ | Normalizing constant for protein motif FAPE | 5.0 |
| A_FAPE,sm_ | …for intra-ligand FAPE | 4.0 |
| A_FAPE,prot-sm_ | …for inter protein-ligand FAPE | 10 |
| A_FAPE,prot(non-motif)_ | …for non-motif FAPE. | 10 |
| D_clamp,prot_ | Clamp upper bound for protein motif FAPE | 5 |
| D_clamp,sm_ | …for intra-ligand FAPE | 4.0 |
| D_clamp,prot-sm_ | …for inter protein-ligand FAPE | 10 |
| D_clamp,prot(non-motif)_ | …for non-motif FAPE | 10 |
| Batch size | Effective batch size on 8GPUs | 16 |
| Learning rate | NA | 0.0005 |
| p_show_motif_seq | Probability of showing the amino acid identity of non-ligand motif tokens | 0.65 |
| Diffusion coordinate scaling | Scaling factor applied to coordinates before the forward process, then inverted and applied after the forward process. | 0.25 |
| Refinement model 3D Gaussian variance | Variance for noising CA atoms in structures. | 1.0 for epochs 1-4, 1.5 for epochs 5-7. |
| Epoch size | Number of examples per training epoch | 25600 |

***Table 4:*** *Training hyperparameters for CA RFdiffusion (diffusion model and refinement model).*

**Loss function**

The loss function to train the diffusion model for CA RFdiffusion is:$L_{diffusion} = W_{disp} L_{disp}+W_{disto,anglo} L_{dist,ang} {+W}_{FAPE,prot} L_{FAPE,prot}+W_{FAPE,prot-sm} L_{FAPE,prot-sm}+W_{FAPE,sm} L_{FAPE,sm}$

Where L_disp_ is the CA coordinate displacement loss (as described in [[1]](https://paperpile.com/c/IXs5yq/m5gK)), L_dist,ang_ is the pairwise inter-residue distogram and anglogram cross entropy loss (as described in [[1]](https://paperpile.com/c/IXs5yq/m5gK)), L_FAPE,prot_ is intra-protein frame-aligned point error (FAPE) [[6]](https://paperpile.com/c/IXs5yq/eX0S) loss on the motif--applied only to pairs of residues which were supposed to be constrained with respect to each other in the motif template definition using Algorithms 1, 2, and 3. L_FAPE,prot-sm_ is the inter-protein-ligand FAPE loss, only computed for frame-point pairs between motif amino acids and ligand atoms (recall, ligand atoms are always considered motif). L_FAPE,sm_ is the FAPE associated with ligands internally only. The values of their respective weights can be found in Table 4.

While L_disp_ and L_disto,anglo_ are identical to how they were defined and applied in [[1]](https://paperpile.com/c/IXs5yq/m5gK), the application of FAPE losses on protein and ligand motif fragments is new to the CA RFdiffusion diffusion model. The FAPE losses were applied to encourage precise and robust reconstruction of ideal motif geometries in the final output of the network. Note that because FAPE by definition scores the quality of an N-CA-C rigid frame orientation, the diffusion model will produce full backbone heavy atom predictions for motif regions, and CA-only predictions in non-motif regions. All small molecules are considered to be a motif and thus encoded in the template structure input.

**Training the refinement model**

Because the diffusion model only produces CA positions in non-motif regions, a second model (“the refinement model”) was trained to take in CA-only descriptions of backbones and produce a refined backbone with all N-CA-C positions and orientations fully described. In short, a model was trained to take in natural protein structures with slightly noised CA positions and random N-CA-C frame orientations, and reconstruct the native backbone heavy atoms. Note this is not a diffusion process, but instead a single shot structure noising and refinement.

**Architecture**

The architecture of this model is precisely identical to that of the diffusion model described above.

**Preparing training examples**

Training dataset, masking strategy and motif templating are very similar to that described above for the diffusion model. The only differences are (1) Instead of noising the resulting structures in a forward diffusion process, a single instance of 3D Gaussian noise is added to all CA atoms and ligand atoms in the structure, and the orientation of N-CA-C frames is randomized. (2) The model is trained in two stages in which the datasets sampled are different: We trained for the first 4 epochs with variance 1.0Å^2^ on the PDB monomer only dataset (i.e., no ligands), then for 3 additional epochs with variance 1.5Å^2^ on both the PDB monomer only dataset and the monomer + ligands dataset. For N-CA-C frame orientation randomization, we simply use the scipy.spatial.transform.Rotation object to produce random rotation matrices for our frames via Rotation.random(L), and orient them accordingly. These examples are then fed to the model, and it is tasked with reproducing the native protein structure.

| **Refinement model training stage** | **Dataset probabilities and mask probabilities** | **3D Gaussian noise variance** |
| --- | --- | --- |
| 1 | Datasets: PDB monomer - 100%  MasksTraining: “Chunked diffusion mask” - 20%  Multi triple contact - 50% | 1.0 |
| 2 | Datasets: PDB monomer - 40%  PDB monomer + ligand - 60%  Masks (for PDB monomer set):  “Chunked diffusion mask” - 20%  Multi triple contact - 50%  Unconditional - 30%  Masks (for PDB monomer + ligand set):  Small molecule contact mask - 100% | 1.5 |

**Table 5:** Dataset proportions, mask sampling proportions, and 3D Gaussian noise variance for the two stage refinement model training procedure.

**Loss function**

The loss function for refinement is very similar to $L_{diffusion}$ defined above:

$$L_{refinement}= L_{diffusion} + W_{FAPE,prot(non-motif)} L_{FAPE,prot(non-motif)}+ W_{pLDDT}L_{pLDDT}$$

Where $L_{FAPE,prot(non-motif)}$is the FAPE loss associated with any non-motif regions (which are always protein only regions), $W_{FAPE,prot(non-motif)}$ is its corresponding weight, $L_{pLDDT}$ is the loss associated with predicting the LDDT of the refined structure (see [[2]](https://paperpile.com/c/IXs5yq/Wd6r) for details of function) with weight factor $W_{pLDDT}$. Additionally, for refinement training we set $W_{disp} =0$ (i.e., no loss from the coordinate displacement loss function) and let FAPE serve as the dominant loss signal. See Table 4 for the weight values for specific components of the loss function.

**Running inference**

Inference is run by running a denoising diffusion trajectory with the trained diffusion model, templating a desired motif and constraining inter-motif-chunk DOFs as the user desires (in this study, we constrain all components of the motif with respect to all others). The output from this denoising trajectory is then fed into the refinement model, which receives a template of the original perfect motif, as opposed to a template of the motif as it was reconstructed by the diffusion model. The refinement model then produces a full heavy atom prediction of the backbone, usually reconstructing the motif to within less than 0.2 Angstroms of the native/input. For sidechains within the motif, we rebuild the rotamers from the input motif onto the corresponding residues in the output, using the sidechain dihedral angles from the original motif, and ideal bond lengths and angles. At this stage, the output is ready for sequence design.

**Algorithm 1: “Chunked diffusion mask”**

The following three functions define how to retrieve a chunked diffusion mask from Table 3.

**```**

```
def _get_diffusion_mask_chunked(xyz, prop_low, prop_high, max_motif_chunks=6):
   """
   Masking strategy that creates discontiguous motifs, possibly unconstrained w.r.t each other.
Parameters:
	xyz (torch.tensor, required): (L,14,3) tensor of coordinates

	prop_low (float, required): lower bound on fraction of protein to mask

	prop_high (float, required): upper bound on fraction of proteins to mask
   """
   chunks, chunk_is_motif, motif_ids, ij, ij_can_see = get_chunked_mask(xyz, prop_low, prop_high, max_motif_chunks)
   chunk_starts = np.cumsum([0] + chunks[:-1])
   chunk_ends   = np.cumsum(chunks)
   # make 1D array designating which chunks are motif
   L = xyz.shape[0]
   mask = torch.zeros(L, L)
   is_motif = torch.zeros(L)
   for i in range(len(chunks)):
       is_motif[chunk_starts[i]:chunk_ends[i]] = chunk_is_motif[i]
   # 2D array designating which chunks can see each other
   for i in range(len(chunks)):
       for j in range(len(chunks)):
           i_is_motif = chunk_is_motif[i]
           j_is_motif = chunk_is_motif[j]
           if (i_is_motif and j_is_motif): # both are motif, so possibly reveal info
               ID_i = motif_ids[i]
               ID_j = motif_ids[j]
               assert ID_i != -1 and ID_j != -1, 'both motif but one has no ID'
               # always reveal self vs self
               if i == j:
                   mask[chunk_starts[i]:chunk_ends[i], chunk_starts[j]:chunk_ends[j]] = 1
               else:
                   # find out of this (i,j) are allowed to see each other
                   ix = tuple(sorted([ID_i, ID_j]))
                   can_see = ij_can_see[ij.index(ix)]
                   if can_see:
                       mask[chunk_starts[i]:chunk_ends[i], chunk_starts[j]:chunk_ends[j]] = 1
   return mask.bool(), is_motif.bool()


def get_chunked_mask(xyz, low_prop, high_prop, max_motif_chunks=8):
   """
   Produces a mask of discontiguous protein chunks that are revealed.
   Also produces a tensor indicating which chunks are given relative geom. info
   Parameters:
   -----------
   xyz (torch.tensor): (L, 14, 3) tensor of atomic coordinates
   low_prop (float): lower bound on proportion of protein that is masked
   high_prop (float): upper bound on proportion of protein that is masked
   """
   L = xyz.shape[0]
   # decide number of chunks
   n_motif_chunks = random.randint(2, max_motif_chunks)
   # decide what proportion of the protein is masked
   # prop cannot result in n_unmasked < n_motif_chunks --> clamp high prop
   max_prop = 1-(n_motif_chunks+1)/L       # add +1 to be safe
   high_prop = min(high_prop, max_prop)
   prop = random.uniform(low_prop, high_prop)
   n_masked   = int(L * prop)
   n_unmasked = L - n_masked
   # decide the length of each chunk by randomly sampling
   # positions to cut a line with n_chunks - 1 cuts
   cuts    = sorted(random.sample(range(1, n_unmasked), n_motif_chunks - 1))
   lengths = [cuts[0]] + [cuts[i] - cuts[i-1] for i in range(1, len(cuts))] + [n_unmasked - cuts[-1]]
   # decide which chunks are given relative geom. info
   # walk over all unique pairs
   motif_pairs = list(itertools.combinations(range(n_motif_chunks), 2))
   # 33% chance that a pair can see each other
   pairs_can_see = [random.choice([True, False, False]) for _ in range(len(motif_pairs))]
   # decide location of chunks within the protein
   # (1) decide order
   random.shuffle(lengths)
   # (2) split available space remaining into other chunks between
   #     the chunks that have been assigned a length
   ngap_low  = len(lengths) - 1
   ngap_high = ngap_low + 1
   ngap = random.randint(ngap_low, ngap_high)
   if ngap == (ngap_low): # there's no cterm/nterm gaps
       Nterm_gap = False
       Cterm_gap = False
   elif ngap == (ngap_low + 1): # there's either a cterm or nterm gap
       Nterm_gap = random.choice([True, False])
       Cterm_gap = not Nterm_gap
   else: # there's both a cterm and nterm gap
       Nterm_gap = True
       Cterm_gap = True
   gaps = sample_gaps(ngap, n_masked) # gaps between unmasked chunks
   random.shuffle(gaps)
   chunks = []
   is_motif = []
   motif_ids = []
   cur_motif_id = 0
   if Nterm_gap:
       chunks.append(gaps.pop())
       is_motif.append(False)
       motif_ids.append(-1)
   for i in range(len(lengths)):
       chunks.append(lengths[i])
       is_motif.append(True)
       motif_ids.append(cur_motif_id)
       cur_motif_id += 1
       if len(gaps) > 1:                       # more to spare
           chunks.append(gaps.pop())
           is_motif.append(False)
           motif_ids.append(-1)
       elif len(gaps) == 1 and not Cterm_gap:  # only one left, but no Cterm gap
           chunks.append(gaps.pop())
           is_motif.append(False)
           motif_ids.append(-1)
       else:
           pass # no more to spare
   if Cterm_gap:
       assert len(gaps) == 1
       chunks.append(gaps.pop())
       is_motif.append(False)
       motif_ids.append(-1)
   assert sum(chunks) == L, f'chunks sum to {sum(chunks)} but should sum to {L}'
   return chunks, is_motif, motif_ids, motif_pairs, pairs_can_see
def sample_gaps(n, M):
   """
   Samples n chunks that sum to M.
      https://stackoverflow.com/questions/2640053/getting-n-random-numbers-whose-sum-is-m/2640079#2640079
   Parameters:
	n (int, required): number of chunks
	M (int, required): number that the chunk lengths should sum to
   """
   nums = np.random.dirichlet(np.ones(n))*M
   # now round to nearest integer, conserving the total sum
   rounded = []
   round_up = True
   for i in range(len(nums)):
       if round_up:
           rounded.append(np.ceil(nums[i]))
           round_up = False
       else:
           rounded.append(np.floor(nums[i]))
           round_up = True
   # ensure all > 0
   for i in range(len(rounded)):
       if rounded[i] < 1:
           rounded[i] = 1
   while sum(rounded) > M:
       ix = np.random.randint(0, len(rounded))
       if rounded[ix] < 2: # must be at least 1
           continue
       rounded[ix] -= 1
   while sum(rounded) != M:
       ix = np.random.randint(0, len(rounded))
       rounded[ix] += 1
   assert all([x >= 1 for x in rounded])
   return [int(x) for x in rounded]
```

**```**

**Algorithm 2: “Multi triple contact”**

The following functions define how to retrieve a “multi triple contact” motif mask

**```**

```
def _get_multi_triple_contact_3template(xyz,
                                        low_prop,
                                        high_prop,
                                        max_triples=2,
                                        xyz_less_than=6,
                                        seq_dist_greater_than=10,
                                        len_low=1,
                                        len_high=7,
                                        force_triples=None):
   """
   Gets 2d mask + 1d is motif for multiple triple contacts.
   Parameters:
       xyz (torch.tensor, required): (L, 14, 3) atomic coordinates
       low_prop (float, required): lower bound for proportion of protein to mask
       high_prop (float, required): upper bound for proportion of protein to mask
       max_triples (int, optional): maximum number of triples to find. Default is 2.
       xyz_less_than (int, optional): maximum distance between atoms to consider a contact. Default is 6.
       seq_dist_greater_than (int, optional): minimum sequence distance between atoms to consider a contact. Default is 10.
       len_low (int, optional): minimum length of motif chunk. Default is 1.
       len_high (int, optional): maximum length of motif chunk. Default is 7.
       force_triples (int, optional): force the number of triples to be this number. Default is None.
   """
   contacts = get_contacts(xyz, xyz_less_than, seq_dist_greater_than)
   if not contacts.any():
       return _get_diffusion_mask_chunked(xyz, low_prop, high_prop, max_motif_chunks=6)
   is_motif_stack = []
   mask_2d_stack = []
   if force_triples is None:
       n_triples = random.randint(1, max_triples)
   else:
       n_triples = force_triples

   for i in range(n_triples):
       indices = find_third_contact(contacts)

       if (indices is None):
           if i == 0:
               # we found no triples at all, so just return a simple chunked diffusion mask
               return _get_diffusion_mask_chunked(xyz, low_prop, high_prop, max_motif_chunks=6)
           else:
               # we found i triples but couldn't find i+1 --> regenerate and return the i triples
               return _get_multi_triple_contact_3template(xyz, low_prop, high_prop, force_triples=i)
       L = xyz.shape[0]
       # 1d tensor describing which residues are motif
       tmp_is_motif = sample_around_contact(L, indices, len_low, len_high)
       # now get the 2d tensor describing which residues can see each other
       # For these, all motif chunks can see each other
       tmp_mask_2d = tmp_is_motif[:, None] * tmp_is_motif[None, :]
       is_motif_stack.append(tmp_is_motif)
       mask_2d_stack.append(tmp_mask_2d)
   is_motif = torch.stack(is_motif_stack, dim=0).bool()
   mask_2d = torch.stack(mask_2d_stack, dim=0).bool()
   is_motif = torch.any(is_motif, dim=0)
   mask_2d = torch.any(mask_2d, dim=0)
   return mask_2d, is_motif
def sample_around_contact(L, indices, len_low, len_high):
   """
   Given a list of indices, sample a revealed motif around each index.
   Parameters:
       L (int, required): length of protein
       indices (list, required): list of indices around which to sample motif
       len_low (int, optional): minimum length of motif.
       len_high (int, optional): maximum length of motif.
   """
   diffusion_mask = torch.zeros(L).bool()
   for anchor in indices:
       mask_length = int(np.floor(random.uniform(len_low, len_high)))
       l = anchor - mask_length // 2
       r = anchor + (mask_length - mask_length//2)
       l = max(0, l)
       r = min(r, L)
       diffusion_mask[l:r] = True
   return diffusion_mask
def get_Cb(xyz):
   """
   Compute Cb given N,Ca,C
   Parameters:
       xyz (torch.tensor, required): shape (batch, L, 3) Cartesian coordinates of atoms
   """
   N  = xyz[...,0,:]
   Ca = xyz[...,1,:]
   C  = xyz[...,2,:]
   b = Ca - N
   c = C - Ca
   a = torch.cross(b, c, dim=-1)
   return -0.58273431*a + 0.56802827*b - 0.54067466*c + Ca
def get_pair_dist(a, b):
   """
   calculate pair distances between two sets of points

   Parameters:
       a,b (torch.tensor, required): shape (batch, L, 3) Cartesian coordinates of atoms
   """
   dist = torch.cdist(a, b, p=2)
   return dist
def get_cb_distogram(xyz):
   Cb = get_Cb(xyz)
   dist = get_pair_dist(Cb, Cb)
   return dist
def get_contacts(xyz, xyz_less_than=5, seq_dist_greater_than=10):
   """
   Computes a 2D boolean mask of contacting residues. True if i,j are contacting, False otherwise.
   """
   L = xyz.shape[0]
   dist = get_cb_distogram(xyz)
   is_close_xyz = dist < xyz_less_than
   idx = torch.ones_like(dist).nonzero()
   seq_dist = torch.abs(torch.arange(L)[None] - torch.arange(L)[:,None])
   is_far_seq = torch.abs(seq_dist) > seq_dist_greater_than
   contacts = is_far_seq * is_close_xyz
   return contacts
def find_third_contact(contacts):
   """
   Finds a third contact for a pair of contacting residues
   Parameters:
       contacts (torch.tensor, required): (L, L) 2d mask of contacts between residues.
   """
   contact_idxs = contacts.nonzero()
   contact_idxs = contact_idxs[torch.randperm(len(contact_idxs))]
   for i,j in contact_idxs:
       if j < i:
           continue
       K = (contacts[i,:] * contacts[j,:]).nonzero()
       if len(K):
           K = K[torch.randperm(len(K))]
           for k in K:
               return torch.tensor([i,j,k])
   return None
```

**```**

**Algorithm 3: Small molecule contact mask generation**

The following function defines how 1D and 2D motif masks were generated for protein-ligand complex examples.

**```**

```
def _get_sm_contact_3template(xyz,
                             is_sm,
                             contact_cut=8,
                             chunk_size_min=1,
                             chunk_size_max=7,
                             min_seq_dist=9):
   """
   Produces mask2d and is_motif for small molecule, possibly with contacting protein chunks revealed.
   Parameters:
       xyz (torch.Tensor): 3D coordinates of protein and small molecule (L, 14, 3)
       is_sm (torch.Tensor): binary tensor indicating which tokens are small molecule atoms (L,)
       contact_cut (float): distance cutoff for contact between protein and small molecule
       chunk_size_min (int): minimum size of revealed chunk
       chunk_size_max (int): maximum size of revealed chunk
       min_seq_dist (int): minimum sequence distance between revealed chunks
   """
   assert len(xyz.shape) == 3
   ca = xyz[~is_sm, 1,:]
   if ca.shape[0] == 0:
       sm_only = True
   else:
       sm_only = False
   sm_xyz  = xyz[is_sm, 1,:]
   dmap = torch.cdist(ca, sm_xyz)
   dmap = dmap < contact_cut
   protein_is_contacting = dmap.any(dim=-1) # which CA's are contacting sm
   where_is_contacting = protein_is_contacting.nonzero().squeeze()
   n_chunk_revealed = random.randint(0,4)
   if (n_chunk_revealed == 0) or (sm_only):
       is_motif = is_sm.clone()
       is_motif_2d = is_motif[:, None] * is_motif[None, :]
       return is_motif_2d, is_motif
   else:
       is_motif = is_sm.clone()
       cur_min_seq_dist = min_seq_dist # could possibly increment this if needed
       for i in range(n_chunk_revealed):
           chunk_size = torch.randint(chunk_size_min, chunk_size_max, size=(1,)).item()
           if len(where_is_contacting.shape) == 0:
               # ensures where_is_contacting is a 1d tensor
               where_is_contacting = where_is_contacting.unsqueeze(0)
           if (where_is_contacting.shape[0] == 0) and (i == 0):
               # no contacts, so sm is only motif
               is_motif = is_sm.clone()
               is_motif_2d = is_motif[:, None] * is_motif[None, :]
               return is_motif_2d, is_motif
           p = torch.ones_like(where_is_contacting)/len(where_is_contacting)
           chosen_idx = p.multinomial(num_samples=1, replacement=False)
           chosen_idx = chosen_idx.item()
           chosen_idx = where_is_contacting[chosen_idx]
           # find min and max indices for revealed chunk
           min_index = max(0, chosen_idx - chunk_size//2)
           max_index = min(protein_is_contacting.numel(), 1+chosen_idx + chunk_size//2)
           # reveal chunk
           is_motif[min_index:max_index] = True
           # update where_is_contacting
           start = max(0,min_index-cur_min_seq_dist)
           end = min(protein_is_contacting.numel(), max_index+cur_min_seq_dist)
           protein_is_contacting[start:end] = False # remove this option from where_is_contacting
           where_is_contacting = protein_is_contacting.nonzero().squeeze()
           if protein_is_contacting.sum() == 0:
               break # can't make any more chunks
       is_motif_2d = is_motif[:, None] * is_motif[None, :] # all tokens can "see" each other
       return is_motif_2d, is_motif
```

**```**

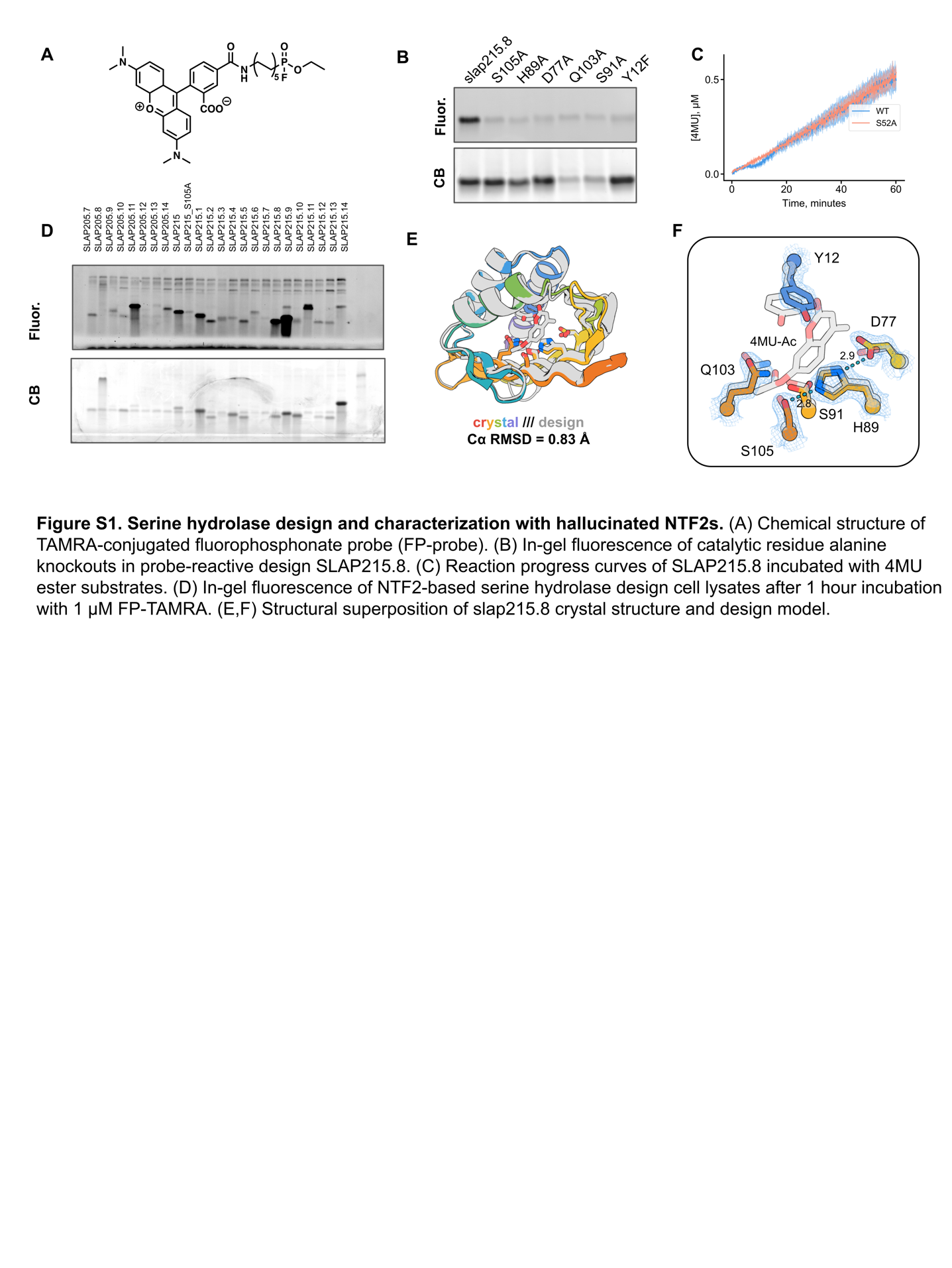

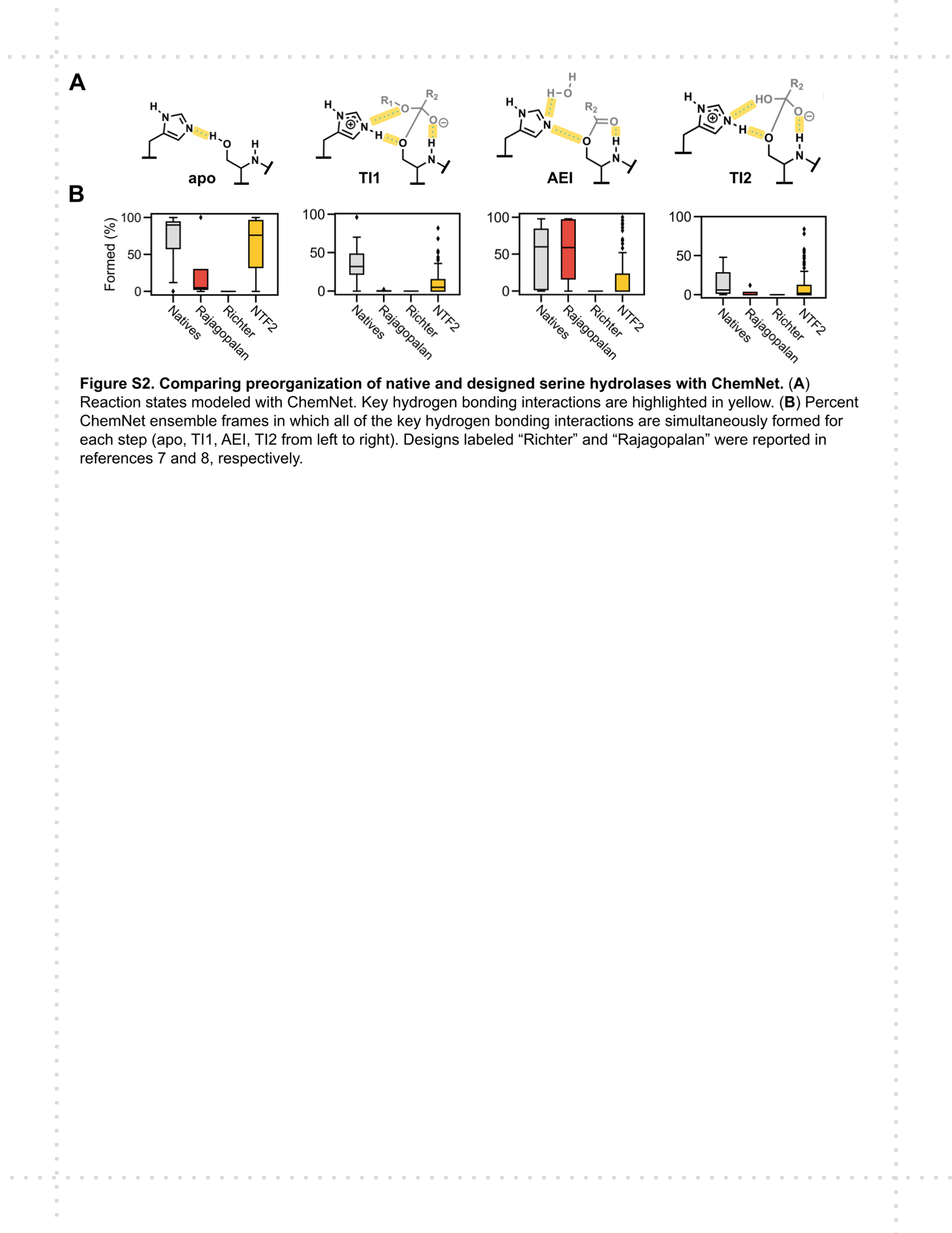

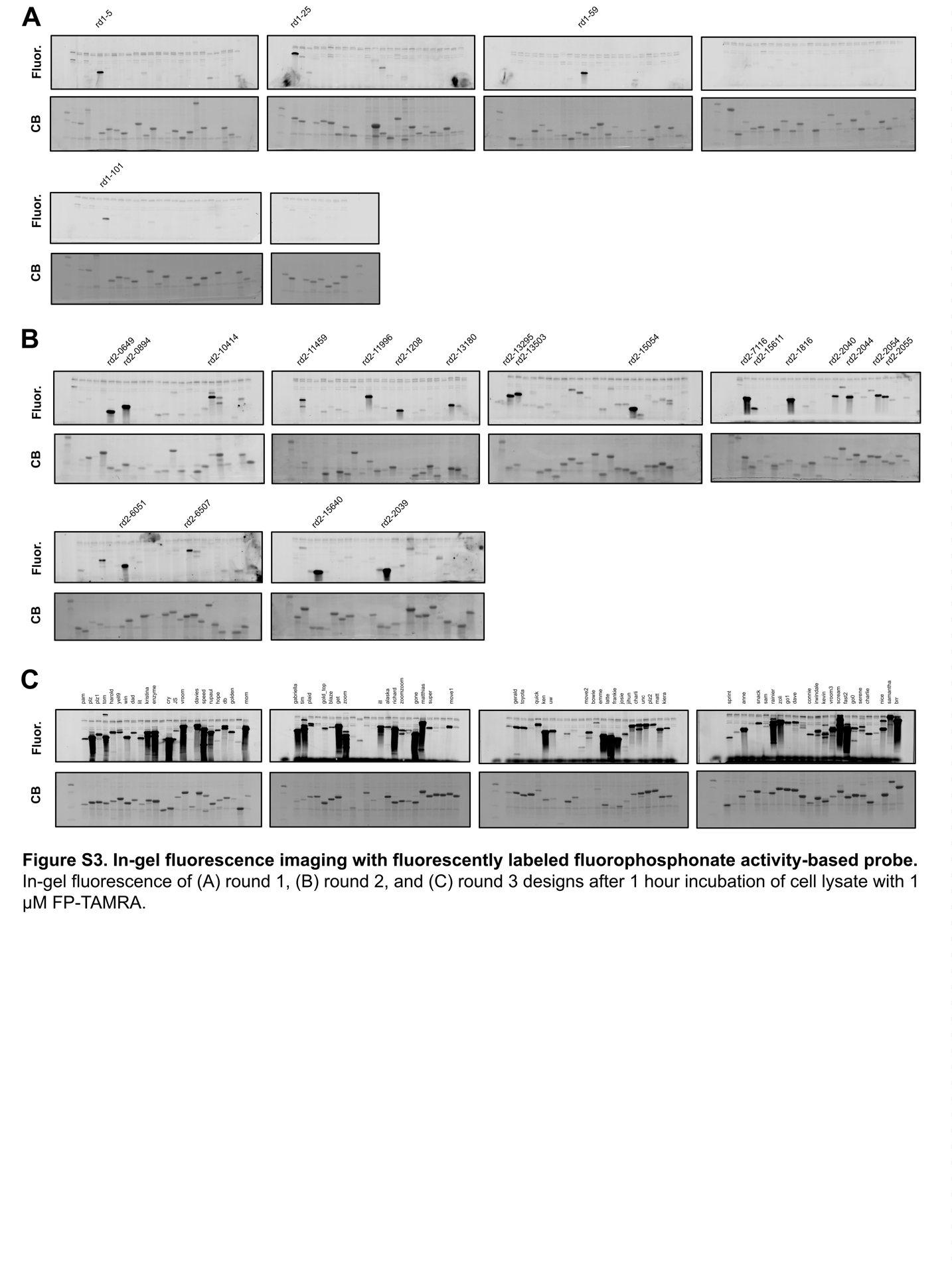

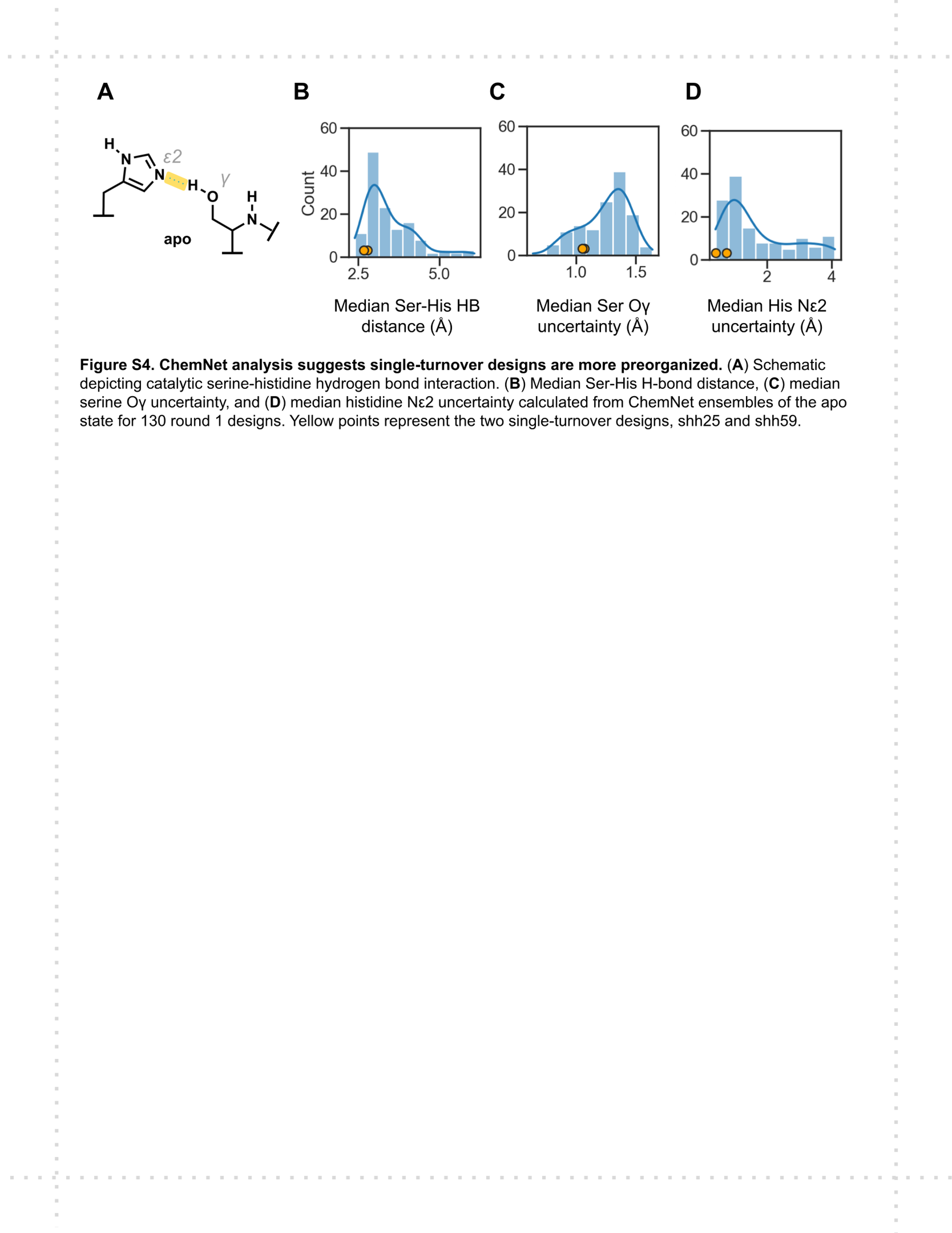

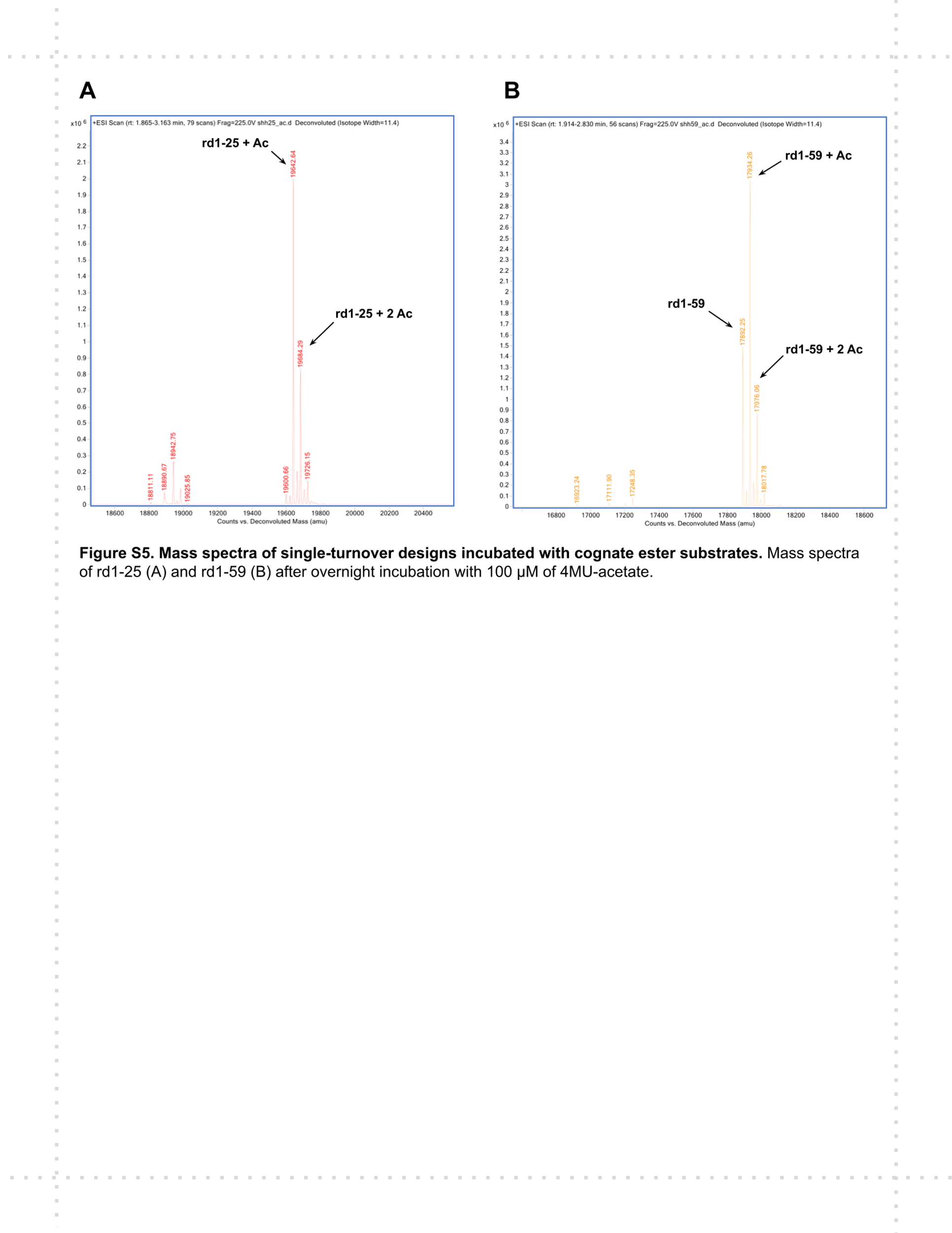

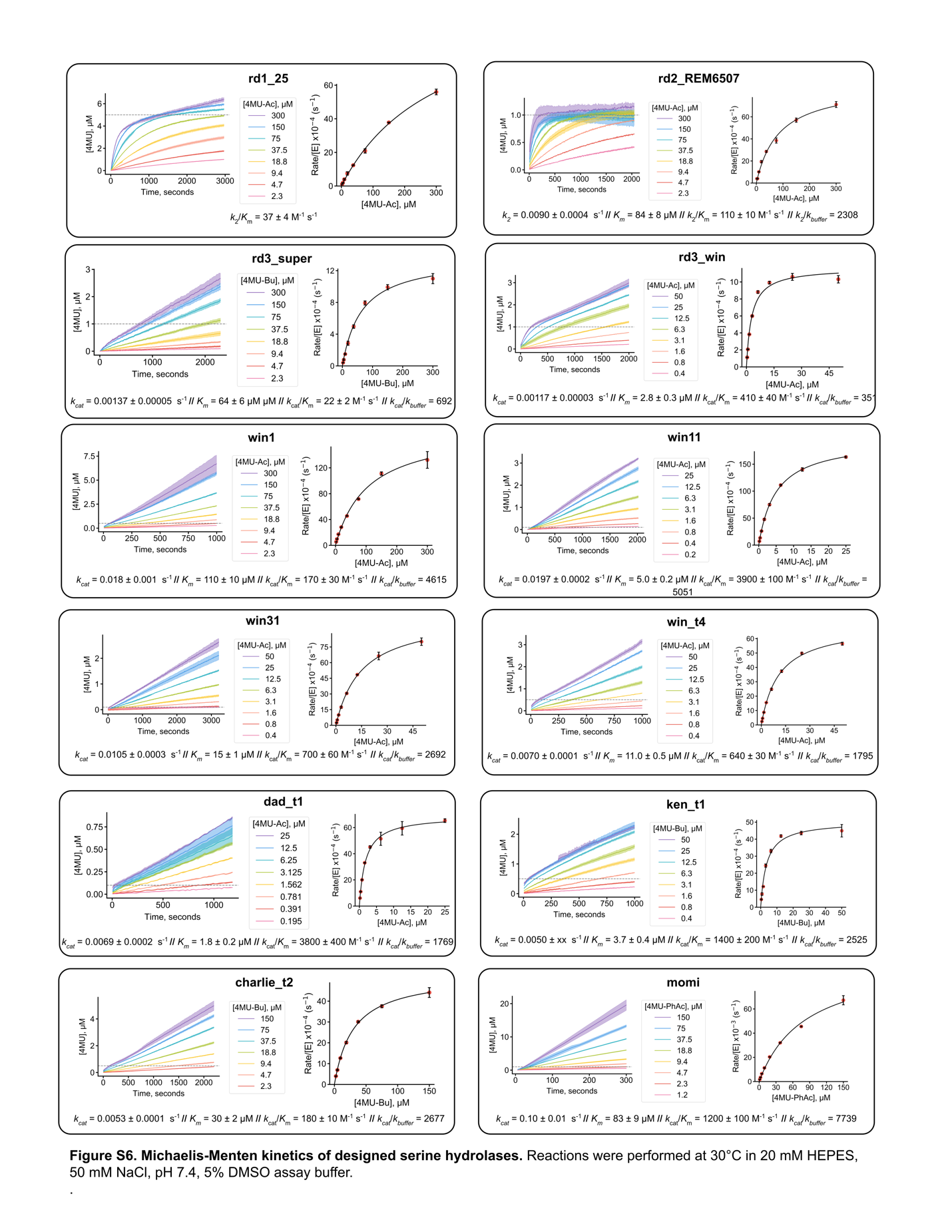

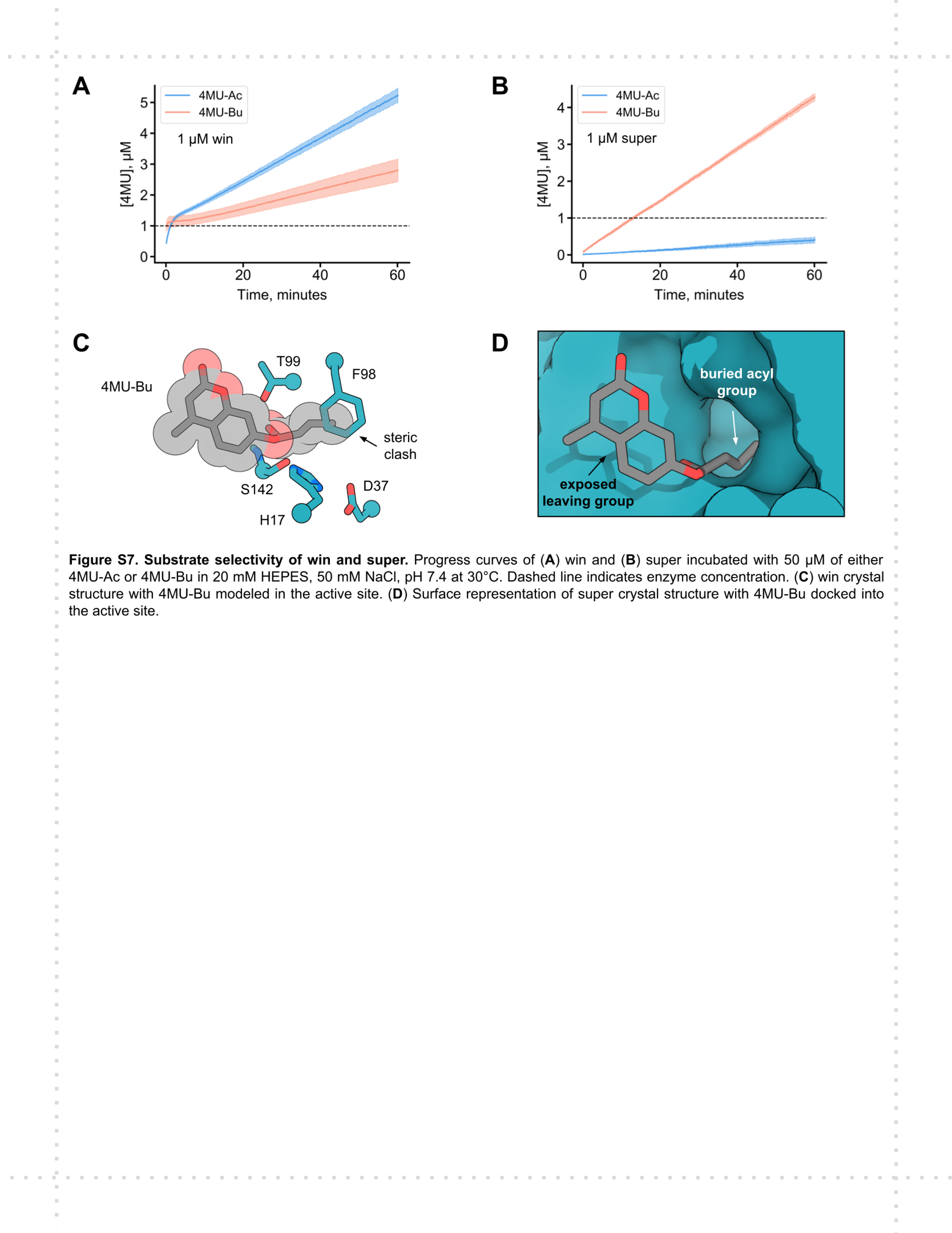

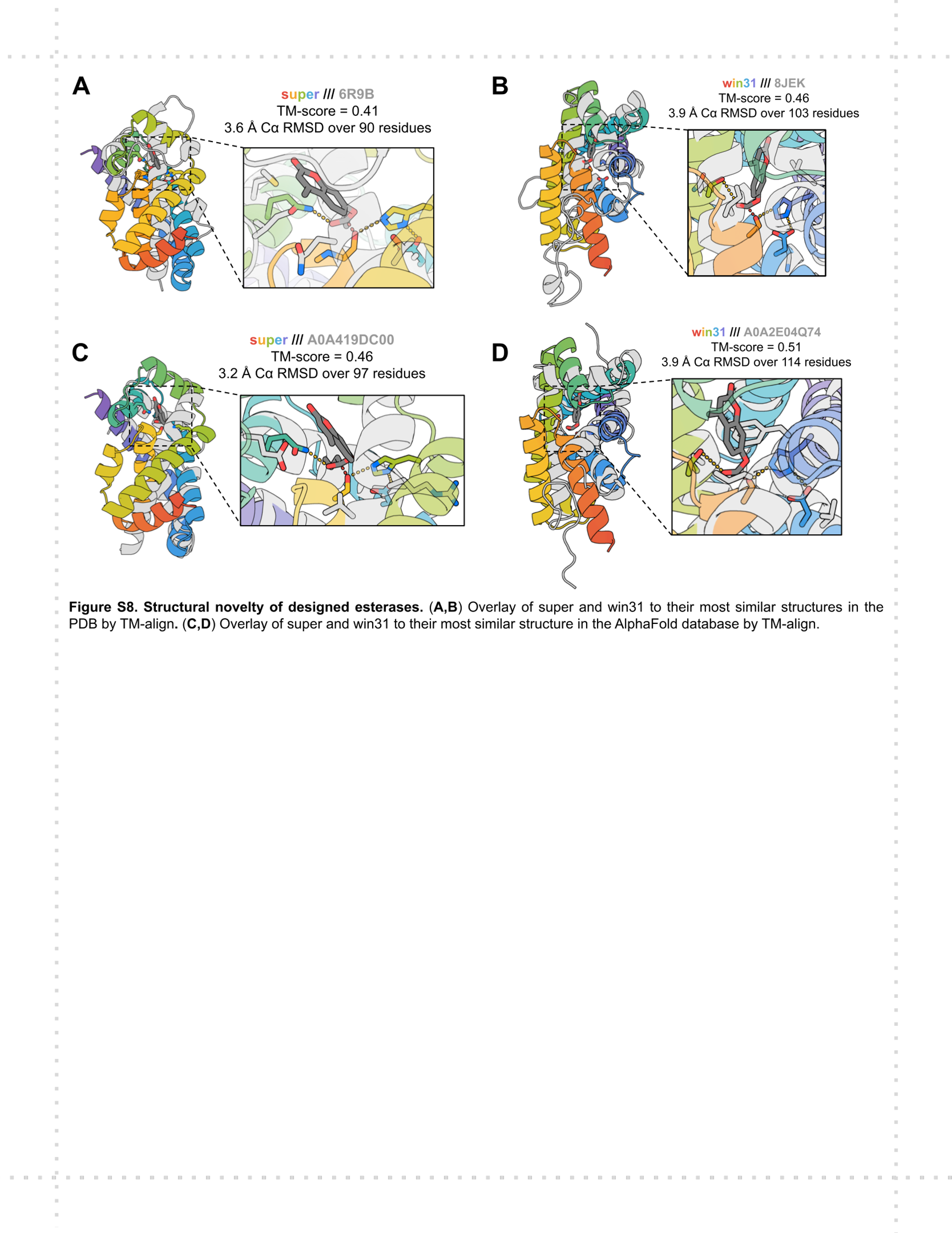

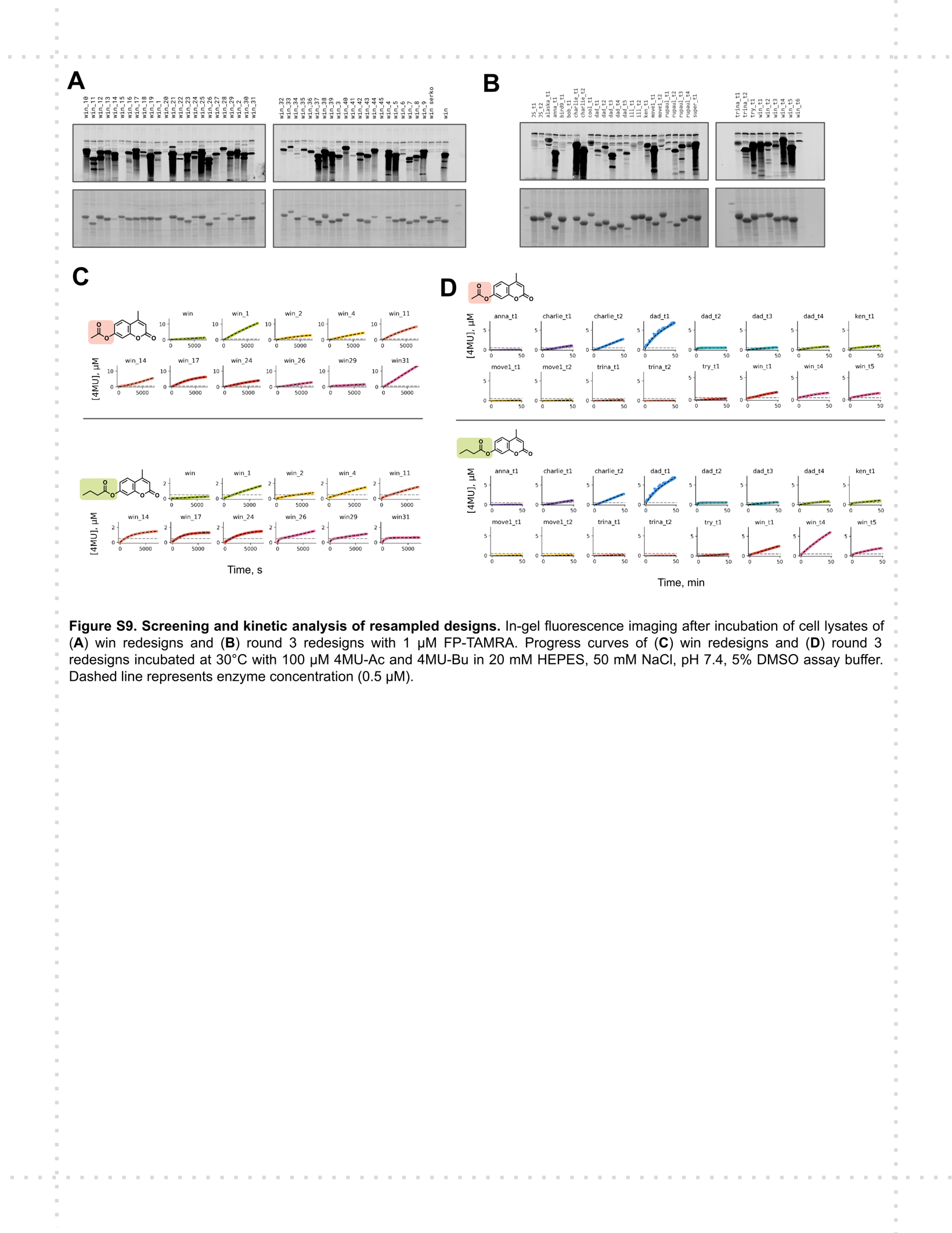

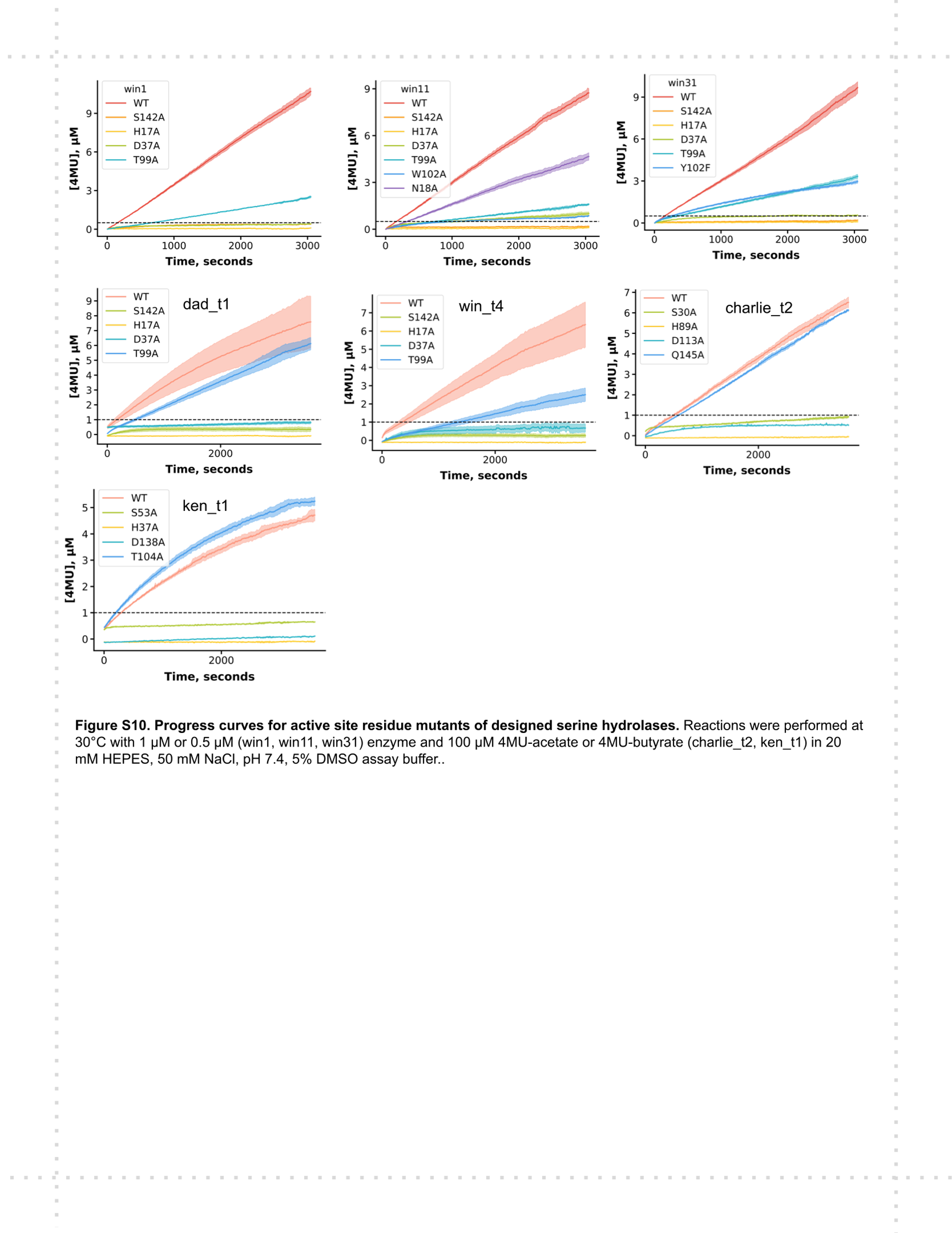

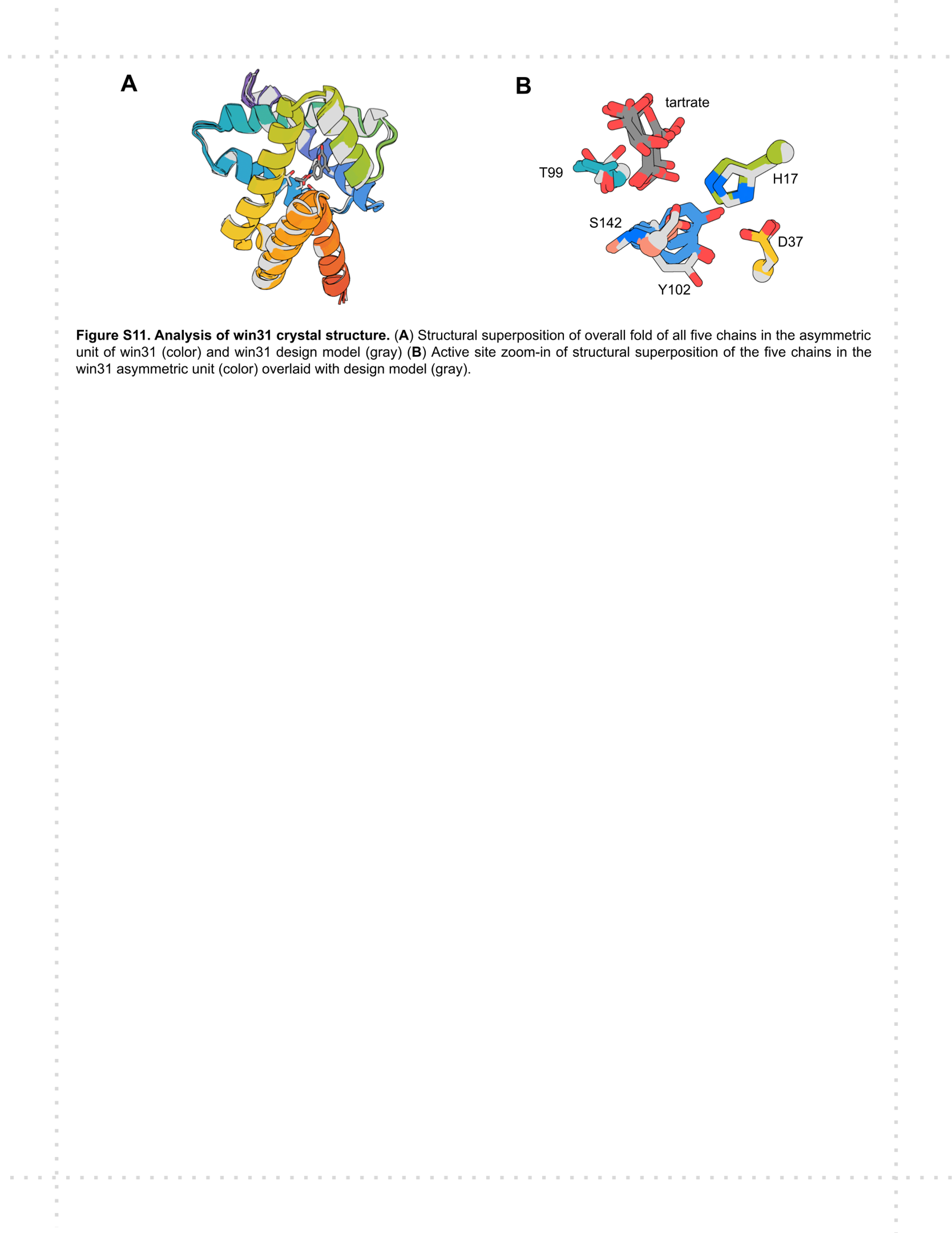

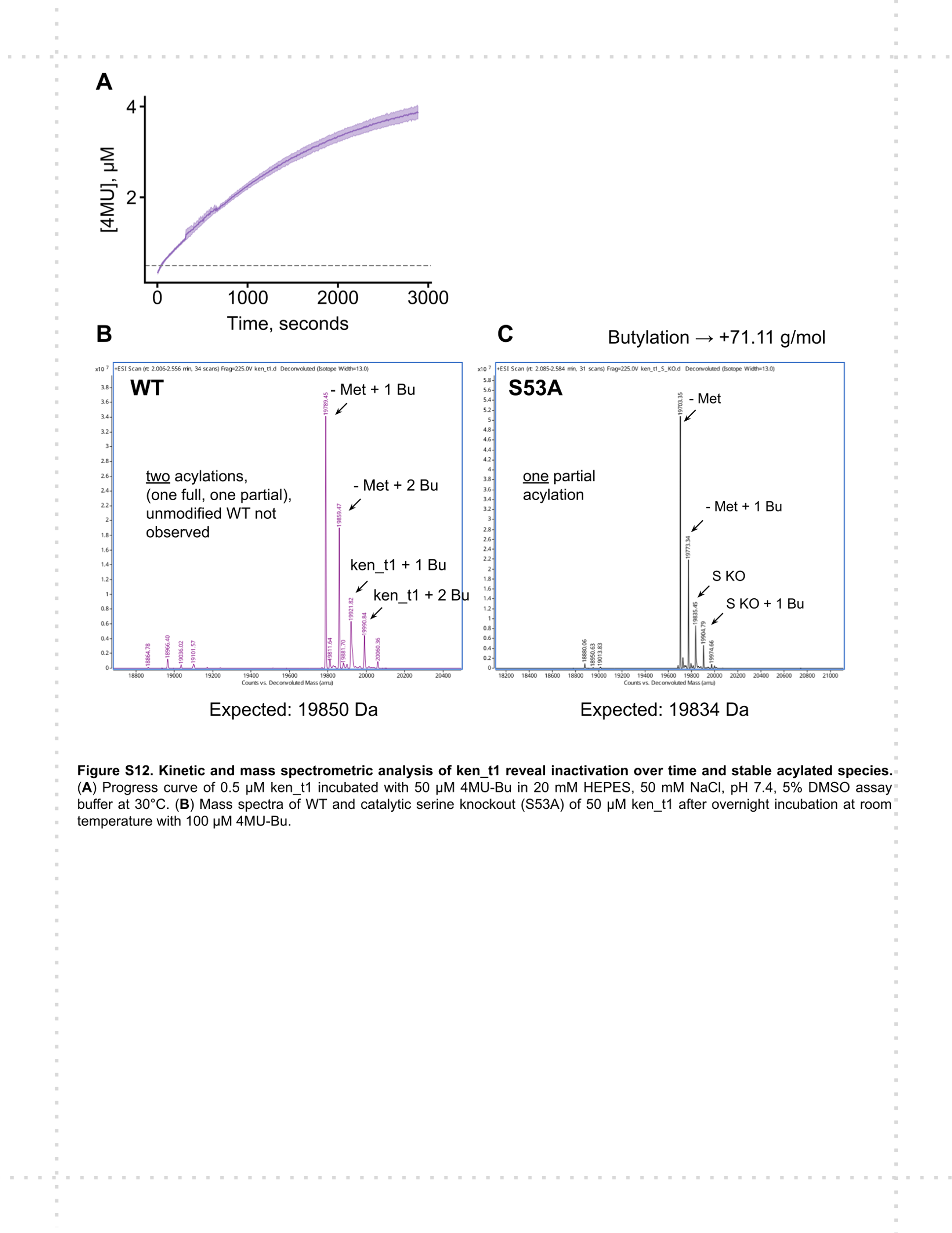

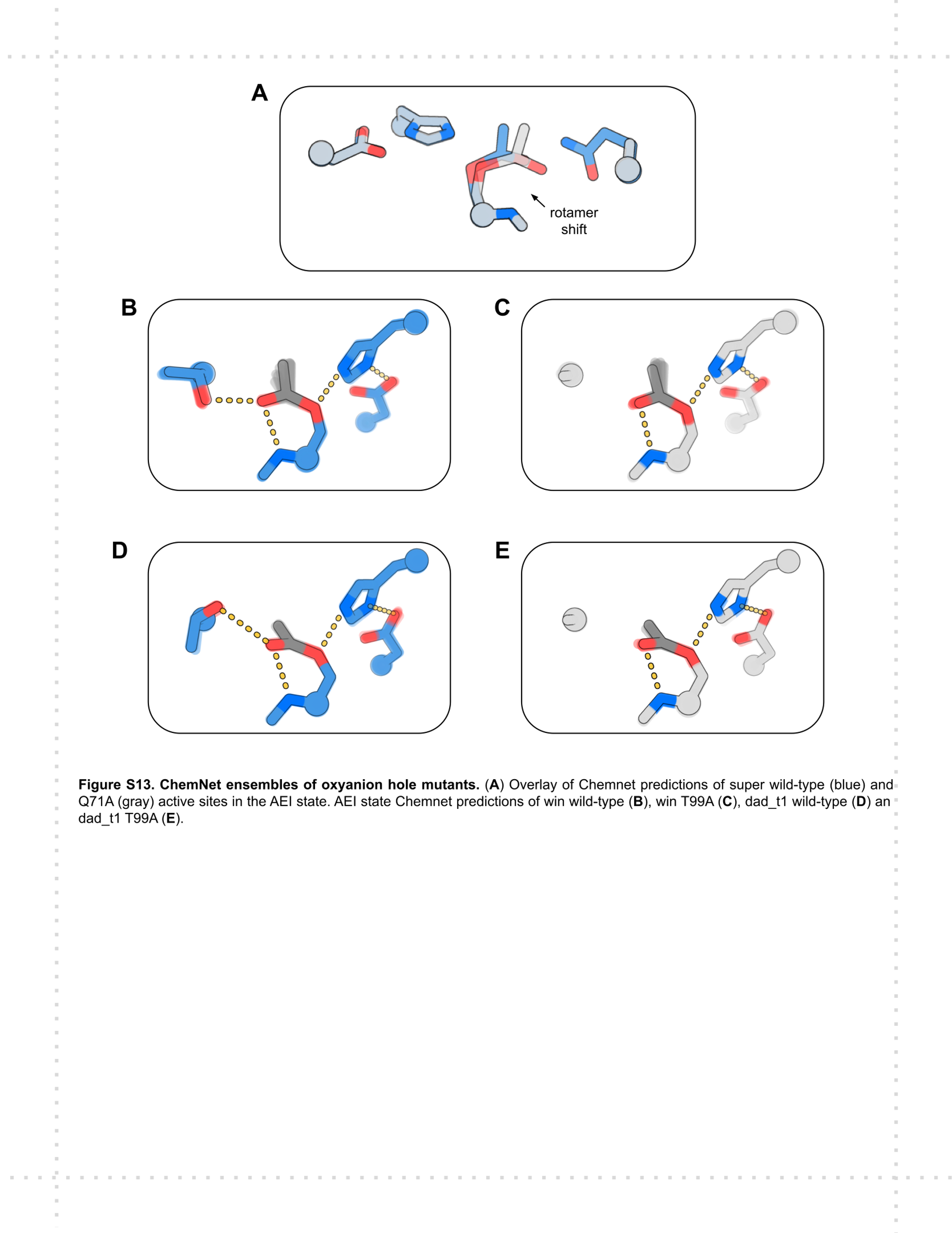

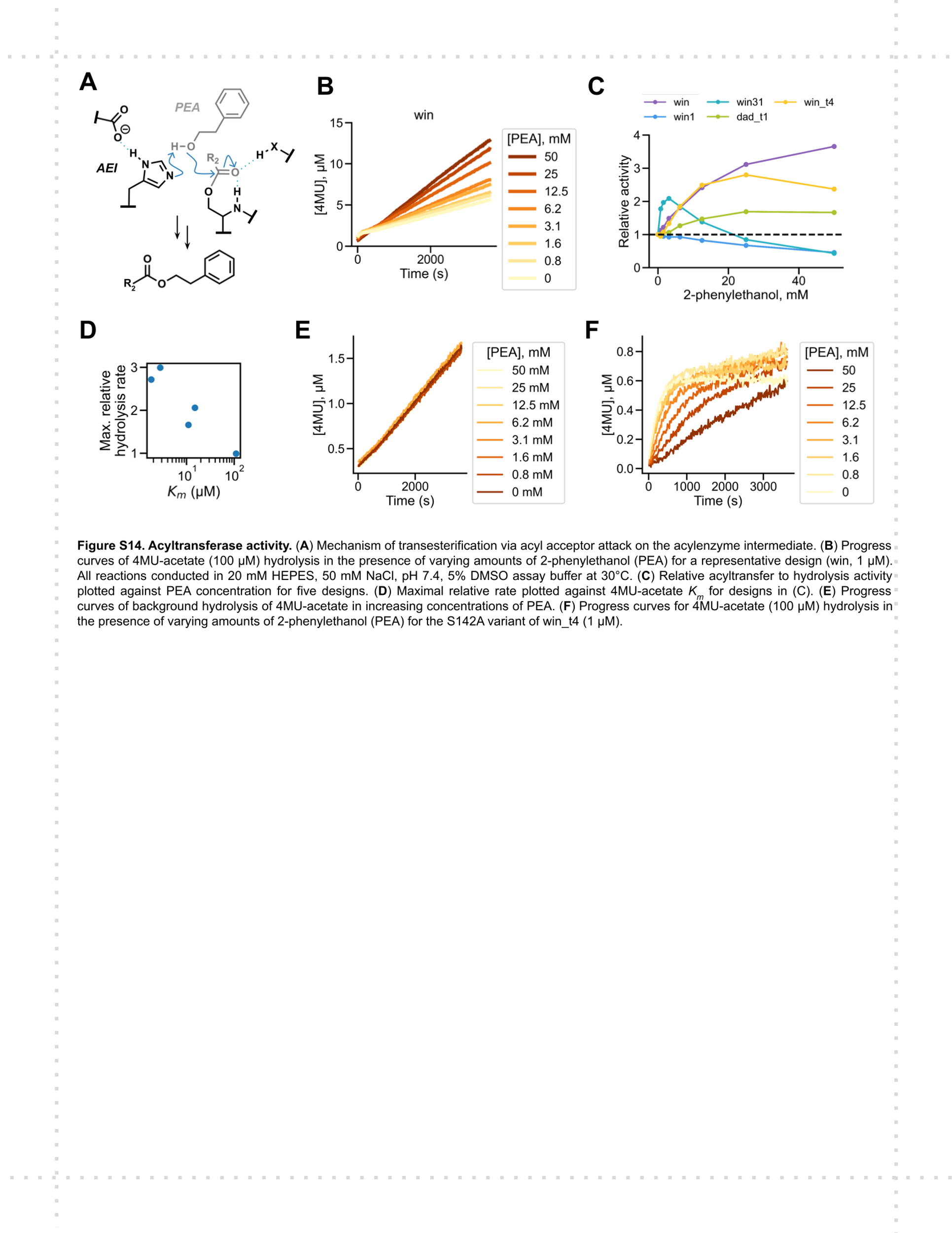

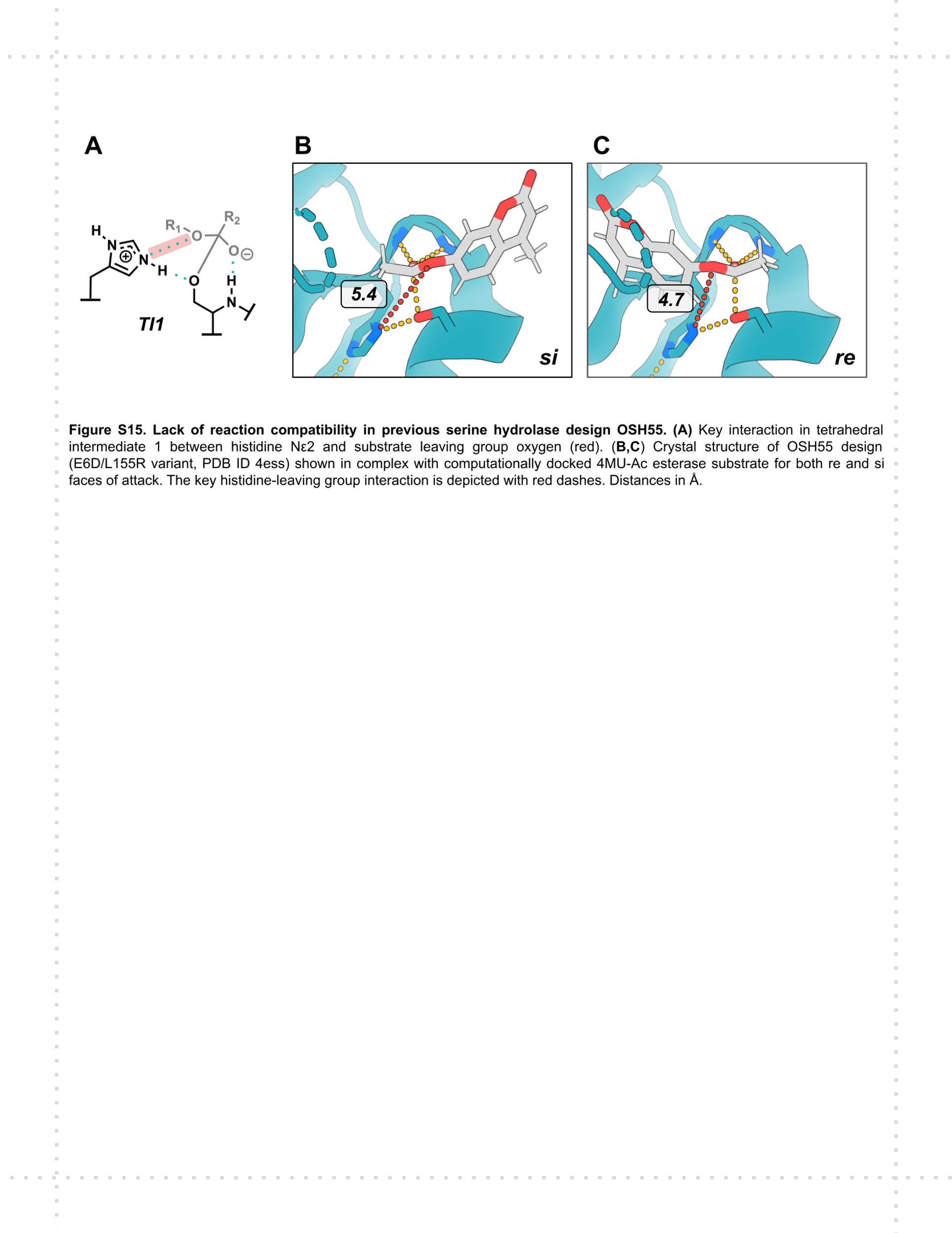
